# Acetylated PLIN2 coordinates the innate immune system and oxylipin synthesis on lipid droplets to propagate inflammation

**DOI:** 10.64898/2026.09.30.755694

**Authors:** Charles P. Najt, Mahima Devarajan, Alex Churchill, Evan Kerr, Lauren Boatner, Samuel Gerrard, Brian Fess, Candace Guerrero, LeeAnn Higgins, Stacy Gelhaus, Douglas G. Mashek

## Abstract

Research has implicated lipid droplets (LDs) and their associated proteins in inflammation and the innate immune response. The key LD protein Perilipin 2 (PLIN2) is tightly correlated with lipid storage and inflammation, but little is known about the mechanisms underlying its proinflammatory effects. Here, we show that PLIN2 is required for the inflammatory response to mitogens and the recruitment of pro-inflammatory proteins to the surface of LDs. PLIN2 directly binds arachidonic acid and interacts with key oxylipin synthetic enzymes such as PLA2 and COX2 to promote LD-localized oxylipin synthesis and propagate inflammatory signaling. Moreover, LPS induces PLIN2 hyperacetylation, and acetylation-dead PLIN2 mutants lose their proinflammatory properties in the presence of an inflammatory stimulus, as if PLIN2 were ablated. Thus, acetylated PLIN2 forms a key metabolon on LDs that is essential for the response and amplification of proinflammatory signaling.

---

Innate immunity is increasingly recognized as a dynamic metabolic system in which lipid handling, organelle homeostasis, and inflammatory signaling are tightly intertwined.^1–3^ Immune cells must continuously adapt their metabolic program to changes in nutrient availability, cellular stress, and microbial or tissue-derived signals.^3^ A central feature of this metabolic–immune interface is the regulated sequestration of neutral lipids within intracellular lipid droplets (LDs), which are now understood as active signaling organelles rather than passive repositories of excess lipid.^1,4^ Dysregulated lipid storage can therefore extend beyond energy balance, shaping the magnitude, duration, and character of innate immune responses.^5,6^

Perilipin 2 (PLIN2) is a major LD-associated protein that links lipid storage to cellular stress responses.^7^ By coating LDs and regulating the accessibility and trafficking of stored lipids, PLIN2 influences lipid accumulation, fatty acid metabolism, and organelle homeostasis across multiple cell types.^8–12^ Notably, PLIN2 is highly responsive to inflammatory and metabolic cues, suggesting that its functions extend beyond the structural maintenance of LDs.^13–15^ In macrophages and other innate immune cells, PLIN2-associated lipid remodeling can influence lipid substrate availability, oxidative stress, mitochondrial function, and inflammatory signaling.^13,16–21^ In hepatocytes, loss of PLIN2 was protective against diet-induced oxidative stress, fibrosis, and inflammatory signaling.^10,12,22^ Conversely, inflammatory stimuli commonly increase PLIN2 expression and LD formation, positioning PLIN2 in a potentially self-reinforcing circuit that connects cellular metabolism to innate immune activation.^1,7,10,23^

Despite growing evidence linking PLIN2 to inflammatory and metabolic disease, fundamental questions remain regarding how PLIN2-dependent LD biology shapes innate immune function. It remains unclear whether PLIN2 primarily serves as a protective adaptation to inflammatory stress, an active driver of immune signaling, or a context-dependent regulator whose effects depend on cellular state and lipid availability. Defining how PLIN2 couples lipid storage to innate immune signaling may thus reveal broader principles governing immunometabolic homeostasis. Here, we examine PLIN2 as a key proinflammatory switch that propagates the immune response. We identify specific PLIN2 acetylation sites that switch PLIN2 into an inflammatory hub, allowing it to coordinate oxylipin biosynthesis at the LD surface and turn LDs into proinflammatory organelles. These results reveal a central role for PLIN2 acetylation in the global inflammatory response and provide insight into the mechanism of LD-mediated inflammation.

## RESULTS

### LDs and the innate immune system

To characterize the impact of acute inflammation on LDs, we injected mice with lipopolysaccharide (LPS) and characterized LDs in the liver, an organ that robustly responds to LPS.^1^ Mice treated with LPS had increased hepatic triglyceride levels and increased LD number (Figure S1A-C). The increase in hepatic triglyceride content corresponded to a decrease in fatty acid oxidation and increased production of oxidized lipid in the form of the inflammatory prostaglandin PGE_2_ (Figure S1D-F). To characterize the enhanced LPS-induced LDs, we fractionated LDs from control and LPS-injected mice (Figure S1G) and performed comparative mass spectrometry profiling of LPS- or control-LD proteins (Figure 1). Analysis of LDs isolated from control and LPS-injected livers identified 4,909 proteins from a total of 61,944 peptides at a 2-peptide and 1.1% false discovery rate (FDR) minimum. LPS-induced LDs showed a significant enrichment of 236 proteins, while only 29 decreased (Figure 1A-B). Pathway analysis mapped protein IDs from the full dataset to pathways largely associated with inflammatory response and metabolism (Figure 1C). The proteins statistically enriched on the surface of LPS-LDs mapped to drug metabolism associated with CYP450 activity, arachidonic acid metabolism, the innate immune response, and DNA adduct metabolism (Figure 1D). Further, 59 upregulated proteins were found only on the LD during LPS treatment (Figure 1B; orange), suggesting the inflammatory response recruits new proteins to the LD surface. These LPS-only LD proteins largely mapped to mitochondrial dysfunction or defense response due to infection and innate immune activation (Figure 1E). Using Kyoto Encyclopedia of Genes and Genomes (KEGG) pathway analysis, statistically significant proteins that clustered together and increased in LPS-LDs mapped broadly to innate immune response and lipid metabolism (Figure 1F; yellow). Specifically, proteins detected only on LPS-LDs clustered around acute inflammatory response, interferon-α response, and cytokine signaling (Figure 1F; solid yellow block). Proteins that were downregulated following LPS clustered around lipase and NAD oxidoreductase activities (Figure 1F; blue), indicating a suppression of fatty acid utilization by mitochondria. This was supported by suppressed β-oxidation in livers of LPS-treated mice (Figure S1D-E). The proteins decreased in these groups included ATGL, PLIN5, and other TAG-hydrolyzing enzymes. We confirmed changes in select proteins in the proteomics dataset and resident LD proteins via Western blot analysis (Figure 1G and Figure S1H-I). Of the three perilipin (PLIN) proteins expressed in the liver, PLIN2 was the only one that significantly increased with LPS (Figure 1G-H) while PLIN3 and PLIN5 decreased (Figure S1H-I). The increase in PLIN2 on the LD surface also correlated with increased eicosanoid-related COX2, cPLA2, and phospho-PLA2 (Figure S1H-I). These data are consistent with the documented impact of LDs in the inflammation cascade.^1^ Together, these data indicate that inflammatory stimulation recruits PLIN2 and a vast array of immune effector proteins to LDs, suggesting LDs become hubs for inflammatory signaling.

**Figure 1.**
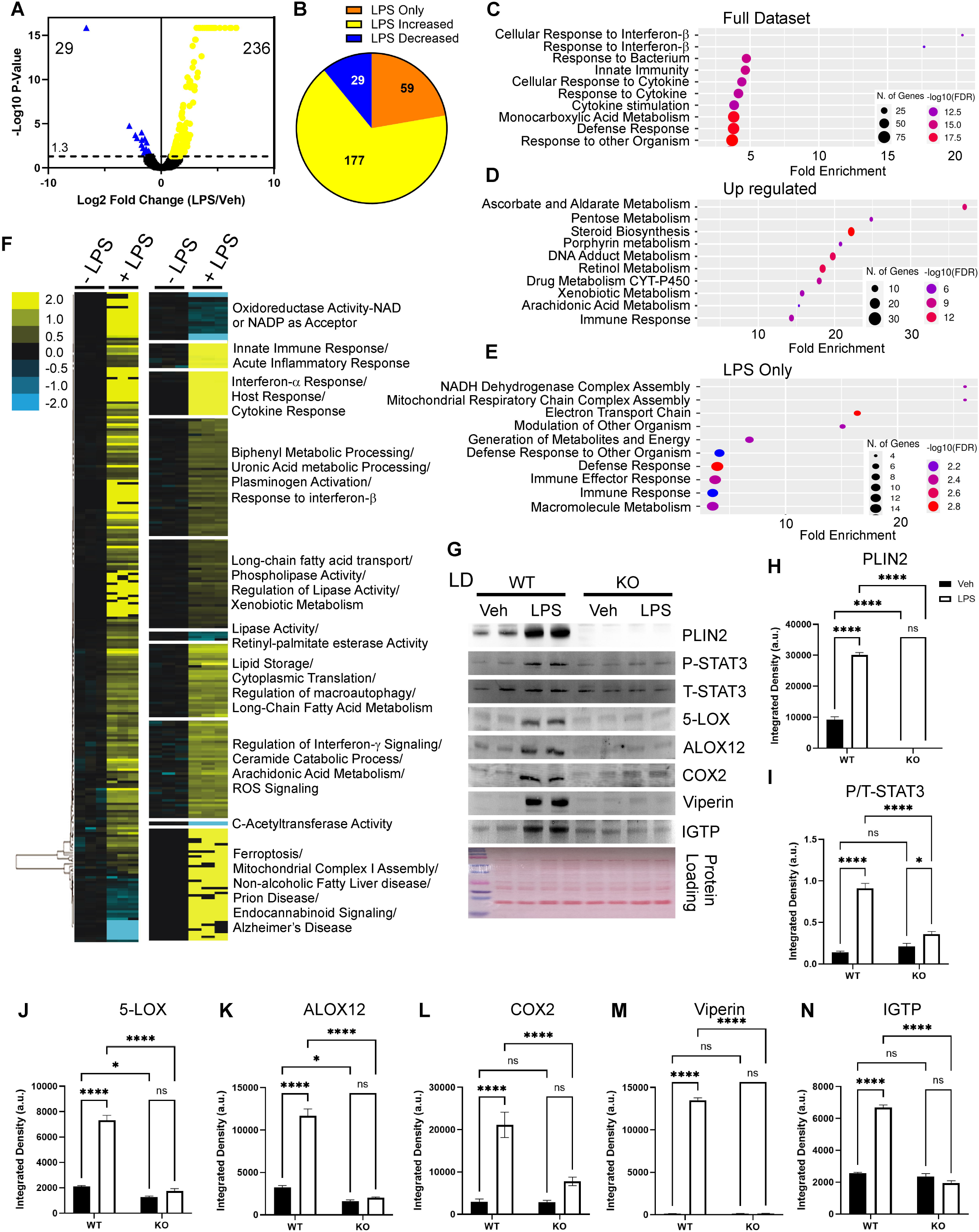
Quantitative mass spectrometry analysis of LDs in response to LPS. A) Volcano plot summarizing the changes in the proteomics of LDs isolated from the livers of control and LPS-treated mice; n =3. Blue indicates proteins that were enriched in the control LD fractions, and yellow indicates LPS-enriched proteins. Black indicates proteins with no significant change due to treatment. Significance was determined using non-parametric quantitative analysis in Proteome Discover. B) Pie chart summarizing the proteomics dataset in terms of the impact of LPS. Orange indicates only proteins present during LPS treatment, yellow indicates proteins that were significantly upregulated due to LPS, and blue indicates proteins that were decreased due to LPS. C) Pathway analysis for all 265 statistically significant proteins. D) Pathway analysis for 236 upregulated proteins that target the LD surface in LPS-treated mice. E) Pathway analysis of the 59 proteins only found on the LD surface of the LDs isolated from mice treated with LPS as compared to control LDs. F) Left: hierarchical clustering of proteins significantly different between control and LPS-induced LDs replicate fractions. The threshold for clustered proteins was determined by the significance between groups. Right: proteins were further grouped using K-means clustering, dividing the significant control and LPS-induced LD proteins into 10 protein clusters. The clustered proteins were mapped to specific metabolic pathways using Panther classification overrepresentation testing. G) LD fractions were analyzed by SDS-PAGE and immunoblotting for resident LD protein PLIN2 and several pro-inflammatory and innate immune proteins. Samples were pooled from two different LD isolations of four animals: control WT and PLIN2 KO mice treated with PBS (Veh) or LPS (5 mg/kg); n=8. Samples were normalized to Ponceau S, and protein abundance was quantified using ImageJ. Significance was determined by two-way ANOVA. n/s indicates no significant difference between the two conditions in the post-hoc test. *P<0.05, ****P<0.01.

### PLIN2 ablation alleviates LPS-induced inflammation

Given recent results^1,10–12,18,24^ and our proteomics screen showing that the innate immune system targets inflammation-induced LDs, we tested whether PLIN2 ablation changes the innate immune response or the recruitment of select proteins to the LD surface (Figure S2 and S3). To test this, we injected wild-type and PLIN2 KO mice (PLIN2 KO demonstrated in Figure S2A-B) with LPS and probed LDs for inflammatory proteins identified in the proteomics dataset. LPS dramatically increased PLIN2, in addition to known anti-pathogenic proteins (STAT3), broad-spectrum anti-microbial peptides (Viperin, IGTP), and oxidized lipid enzymes associated with the arachidonic acid signaling cascade (COX2, ALOX12, ALOX5) (Figure 1G-N). Loss of PLIN2 universally blocked the accumulation of these proteins at the LD surface (Figure 1G-N). In addition to recruiting innate immune proteins to the LD, PLIN2 was essential for upregulating gene transcripts associated with the inflammatory response (Figure S2B-P). While *Ptgs2* expression was suppressed by PLIN2 KO, *Ptgs1 (Cox1)* was refractory to PLIN2 KO mice (Figure S2N), indicating that PLIN2 affects specific arms of the immune response without universal blockade. We also examined 111cytokines and innate immune proteins in the livers of mice using a proteomic profiling assay and observed, similar to the gene expression data, that inflammatory proteins in WT livers were upregulated after LPS injection in a PLIN2-dependent manner (Figure S3 A-B). Taken together, PLIN2 ablation protected against the innate immune response, suppressing both transcription and the recruitment of select proteins to the LD surface.

### PLIN2-coated LDs are the site of oxylipin production

Given that LDs accumulate with LPS stimulation, recruit pro-inflammatory machinery to their surface, and that PLIN2 ablation negates the inflammatory response, we tested whether PLIN2 localizes inflammatory molecule biosynthesis to LDs. Treatment with LPS increased LDs in AML12 cells (Figure S4A-B), in addition to increasing phospholipase activity from phospholipids (Figure S4C) and the proinflammatory arachidonic acid metabolite PGE_2_ (Figure S4D). We confirmed this was not unique to immortalized cells - primary hepatocytes isolated from WT mice treated with vehicle or LPS similarly increased LDs and PGE_2_ levels (Figures S4E-G). To visualize where this oxylipin synthesis occurred, we treated AML12 cells with CoxFluor, a fluorescent sensor that detects COX2 activity in real time^25^, and counterstained for LDs. These studies showed fluorescent activity in the cytosol and puncta at or near the LD surface (Figure 2A; top set of images). When the cells were treated with LPS, the CoxFluor signal increased 7-fold relative to the baseline PBS-treated cells (Figure 2A and B; bottom set of images). Moreover, the CoxFluor signal shifted from weak and diffuse in the cytosol to a strong signal at or near the LD surface (Figure 2B; bottom set of images). The correlation of the CoxFluor sensor with the LD stain increased 3-fold with LPS, indicating that COX2 activity and oxylipin production occur at the LD surface (Figure 2B-C). We repeated these studies in eGFP-PLIN2-transfected cells (Figure 2D-F). Under inflammatory conditions, the CoxFluor signal again increased ∼7-fold relative to PBS-treated cells, while the pattern of the CoxFluor signal was the most intense at or near PLIN2 (Figure 2D). PLIN2 co-localized with 82% of the CoxFluor signal (Figure 2E), suggesting that PLIN2 is associated with COX2 oxylipin production. Moreover, cross-correlation analysis showed that the COXFluor signal overlapped with the eGFP-PLIN2 signal at the edges of the neutral lipid stain (Figure 2F). Using the EicosaCell assay, in which the oxylipin PGE2 is crosslinked to proteins at or near the site of synthesis,^26^ we also recapitulated the COXFluor results in primary hepatocytes by demonstrating enriched PGE_2_ production at PLIN2-coated LDs (Figure S4H).

**Figure 2.**
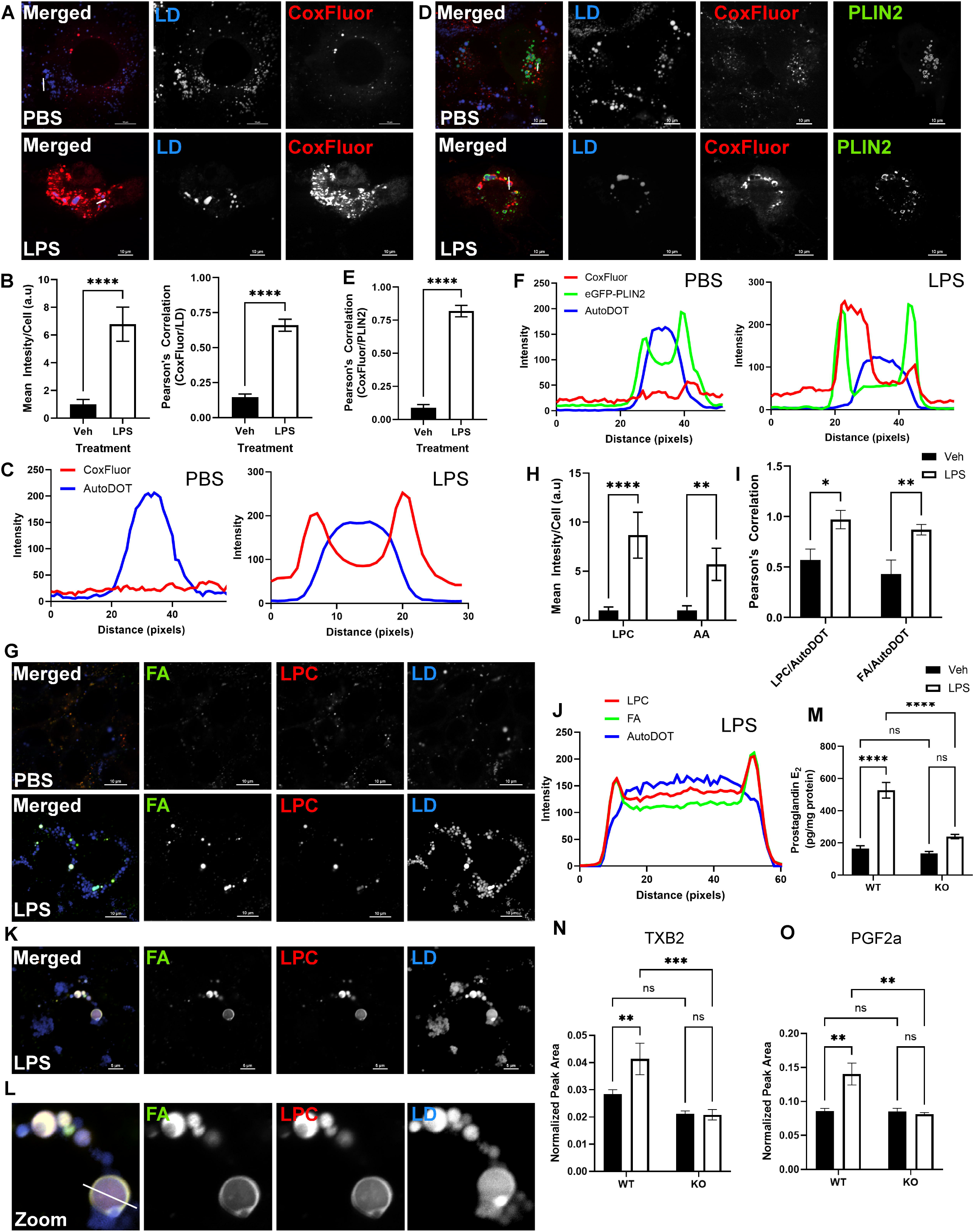
LPS-induced PLIN2-LDs are the site of oxylipin production. A) Representative confocal microscopy images of LDs in AML12 cells in control and LPS medium. Cells were treated with 1 μg/ml LPS for 16 h. Cells were stained with AutoDOT and the COX2 activity sensor CoxFluor. B) Quantification of the CoxFluor intensity within control and LPS-treated cells in (B-C); n=8-12 cells per condition. B) Pearson’s correlation was used to determine interactions of the CoxFluor-COX2 sensor and AutoDOT staining of LDs. C) Cross-correlation indicates that COX2 enzymatic activity is happening at the interaction between the cytosol and the neutral core as indicated by the AutoDOT staining. D) Representative confocal microscopy images of AML12 cells transfected with eGFP-PLIN2. Cells were treated similarly to (A); n=4-6 images with 12-16 cells per condition. E) Colocalization analysis of the interaction between the CoxFluor-COX2 sensor and eGFP-PLIN2. F) Cross-correlation indicates that COX2 enzymatic activity is happening at the interaction between the cytosol and the neutral core, as indicated by the AutoDOT staining where it overlaps with the eGFP-PLIN2 signal. G) Red/Green BODIPY PC-A2 probe is hydrolyzed by PLA_2_ to release a red, fluorescent BODIPY-labeled LPC (red LPC) and a green fluorescent BODIPY linked to FA (Green FA); the site of PLA_2_ activity during LPS treatment was determined. Cells were loaded with the PC-A2 probe, then treated with Veh or LPS for 6-12 h. Confocal images representing the site of PLA_2_ activity; n=3-4 images with 6-12 cells per image. H) Quantification of the two fluorescent BODIPY molecules. I) Colocalization analysis of the interaction between the LPC channel and the neutral lipid AutoDOT stain, as well as the FA channel and the neutral lipid AutoDOT stain. J) Cross-correlation indicates that both the LPC and the FA are at or near the LD surface, as indicated by the overlapping LPC and FA peaks just at the intersection of the AutoDOT counterstain. K-L) Select images focusing on PC-A2 PLA_2_ activity at the LD surface. L) Zoom highlighting the ring-like structure of the PC-A2 PLA2 activity sensor surrounding the neutral lipid core of LDs as indicated by the AutoDOT counterstain. M-O) Oxylipin species produced by COX2 metabolism of arachidonic acid were measured as described in the methods. M) PGE_2_ was quantified via ELISA from livers of control WT and PLIN2 KO mice treated with PBS-Veh or LPS. n=4-6 mice per group. N) TXB2 and O) PGF_2α_ were quantified via LC/MS/MS from livers of control WT and PLIN2 KO mice treated with PBS-Veh or LPS. n=4-6 mice per group. Significance was determined by Student’s T-test for imaging experiments. Significance was determined by two-way ANOVA for oxylipin measurements. *P<0.05, **P<0.01, ***P<0.005, ****P<0.001.

Oxylipins are a broad class of bioactive metabolites formed from the enzymatic and non-enzymatic oxidation of fatty acids (FA). Well-studied oxylipin subclasses include eicosanoids, prostaglandins (PGs), thromboxanes, and leukotrienes, which are derived from arachidonic acid and docosahexaenoic acid. The lipase responsible for releasing arachidonic acid and docosahexaenoic acid for enzymatic and non-enzymatic production of oxylipins is phospholipase A2 (PLA_2_; *Pla2g2a*). Several studies have implicated PLA_2_ enzymatic activity at or near the LD surface during LD biogenesis^27–29^ and viral replication^28,30^, but its role in LD-associated inflammation is unclear. To address this, we used a fluorogenic phosphatidylcholine with quenched fluorophores attached to each of the two fatty acids, PC-A2. This sensor is specific to PLA_2_ lipolytic activity, as PLA_2_-mediated cleavage of the fatty acid at the sn-2 position of the fluorogenic substrate allows for the subcellular localization of the resulting LPC, and the free fatty acid can be tracked in real time by confocal microscopy (Figure 2G). Under basal conditions, the PC-A2 PLA2 sensor signal was minimal (Figure 2G; top panel). LPS-induced inflammation dramatically increased the PC-A2 PLA_2_ sensor signal around the LD (Figure 2G; bottom panel). The sensor signal increased in both the free-fatty acid (green) and LPC (lysophosphatidylcholine - red) channels (Figure 2H). Increased PLA_2_ activity resulted in a 9.6-fold increase in LPC and a 6.2-fold increase in fatty acid (Figure 2H). We observed the PC-A2 signal pattern at the LD monolayer under LPS conditions (Figure 2I-J). Both the LPC and fatty acid signals surrounded the LD, indicating that PLA2 acted on the LD monolayer and released fatty acids at the LD surface (Figure 2K-L; zoom). Together, these data indicate that pro-inflammatory oxylipin enzymes target PLIN2-coated LDs during inflammatory conditions, using the phospholipid monolayer to generate substrates for oxylipin production.

### PLIN2 ablation blocks pro-inflammatory and increases anti-inflammatory, pro-survival oxylipin production

Given data indicating that oxylipins are produced at the surface of PLIN2-coated LDs, we tested whether PLIN2 ablation alters oxylipin levels during inflammation. To test this *in vivo*, we injected WT and PLIN2 KO mice with LPS and measured hepatic oxylipins by ELISA or LC/MS (Figure 2M-O and S5-6). We detected many oxylipins in hepatic tissue, with several showing expected elevations in response to LPS. Oxylipins are broadly categorized into lipids generated by cyclooxygenases (COX), lipoxygenases (LOX), or cytochrome P450S (CYPs). LPS stimulated production of the COX products PGE_2_, TXB2, and PGF2α in WT mice, while ablation of PLIN2 suppressed their accumulation (Figure 2M-O, S5I-K). We also measured oxylipin levels produced by LOX and CYPs (Figure S5-6). For the pro-inflammatory LOX (Figure S5) and CYP (Figure S6) metabolites, LPS increased levels of 5-HETE and 15-HETE in WT mice, while ablation of PLIN2 blocked the increase (Figure S5A and F). The lack of 5-HETE and 15-HETE in LPS-treated PLIN2 KO mice relative to their controls was consistent with our observation of reduced ALOX5 and ALOX15 expression (Figure S2I and J). 20-HETE was elevated, and 11-HETE was decreased in both WT and PLIN2 KO mice (Figure S5D and G). In contrast, LPS increased 9-HETE and 12-HETE levels in PLIN2 KO mice but had no effect in WT animals (Figure S5C and E). 20-HETE can further be metabolized to form broad chemotaxis mediators 5,15-DiHETE, 8,15-DiHETE, 5-oxoETE, and 12-oxoETE.^31^ LPS decreased 5,15-DiHETE, and 8,15-DiHETE in both WT and PLIN2 KO mice (Figure S5U-V), while increasing 5-oxoETE, 12-oxoETE, and15-oxoETE in only the PLIN2 KO mice (Figure S6O-Q). These three metabolites suppress natural killer cell maturation and NFκB signaling.^32,33^ Neither metabolite responded to LPS in control animals, but PLIN2 ablation increased both metabolites (Figure S6O-Q). The remaining two 20-HETE metabolites we detected were 14,15-DiHETE (Figure S5W). In support of PLIN2 ablation enhancing anti-inflammatory pathways, PLIN2 KO mice had elevated levels of the anti-inflammatory pro-resolving oxylipins 11-HEPE, 12-HEPE, 15-HEPE, 11,12-DiHET, and 14,15-DiHET (Figure S5Q-S and S6F). In summary, while LPS commonly elevated proinflammatory oxylipins, PLIN2 ablation blunted their production. This correlates with accumulation of anti-inflammatory, pro-cell-survival oxylipins derived from other fatty acid species.

### PLIN2 directly interacts with innate immune and oxylipin enzymes at the LD surface

Given PLIN2’s role in oxylipin production and recruitment of inflammatory proteins to the LD surface, we used quantitative immunoprecipitation^34^ followed by LC-MS/MS to characterize the PLIN2 interactome under control and LPS conditions. Analysis of PLIN2-interacting proteins identified 2,490 proteins from a total of 24,961 peptides at a 2-peptide minimum and a 0.8% FDR. LPS-induced PLIN2 interactions showed a significant enrichment of 236 proteins (Figure 3A-B). Pathway analysis mapped the 236 LPS-driven PLIN2 interactions primarily to numerous metabolic pathways and metabolic and neurodegenerative diseases. In contrast, LPS decreased the expression of only 11 proteins that interacted with PLIN2 (Figure 3A-B). These proteins were highly specific and mapped to cytoskeletal binding and locomotion, as well as cytosolic DNA sensing related to MAPK/ERK signaling. PLIN2 interacted with 145 proteins only under LPS conditions (Figure 3B). These genes mapped to pathways associated with oxidative phosphorylation, ROS signaling, and arachidonic acid signaling, as well as several metabolic and age-related diseases (Figure 3C). Hierarchical clustering further highlighted specific sets of proteins differentially enriched in control and LPS-induced PLIN2 interactions, beyond global pathways (Figure 3D). Using K-means clustering and KEGG pathway analysis, statistically significant proteins that clustered together and were increased in LPS-PLIN2 interactions mapped to largely to inflammatory signaling (Figure 3D; yellow). Genes downregulated by LPS clustered around longevity-regulating pathways, including actin and cytoskeleton organization, DNA damage response, and insulin signaling (Figure 3D; blue). Figure 3E presents a schematic of proteins PLIN2 interacts with. In this diagram, PLIN2 was linked to upstream regulators known to activate the inflammatory cascade, including HMOX1, TIFA, and Viperin. Additionally, PLIN2 interacted with PLA2G4A, PLA2G7, and FAAH, enzymes known to release and regulate arachidonic acid levels. Downstream of the phospholipases, PLIN2 was also shown to interact with several CYPs, LOX, and eicosanoid-producing enzymes (ALOX12, PTGES2, Cyp4a12, Cyp2e1). These results support PLIN2’s role in the innate immune response and the arachidonic acid signaling cascade, linking our findings of blunted innate immune genes and proteins to changes in phospholipid and oxylipin levels.

**Figure 3.**
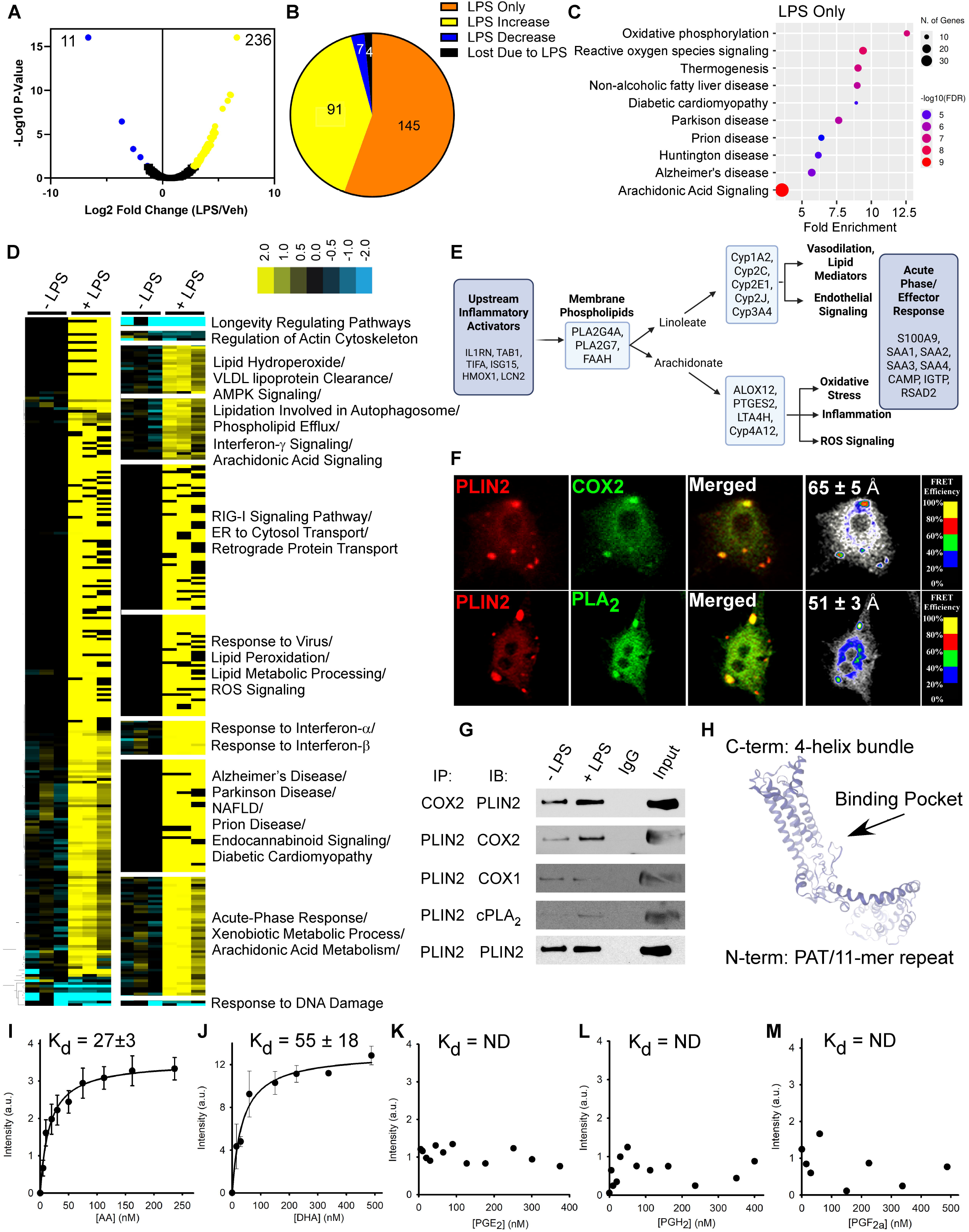
PLIN2 directly interacts with innate immune and oxylipin-producing enzymes. A) Volcano plot summarizing the changes in the proteomics of quantitative PLIN2 immunoprecipitation from the livers of control and LPS-treated mice; n =3. Blue indicates proteins that were enriched in the control LD fractions, and yellow indicates LPS-enriched proteins. Significance was determined using non-parametric quantitative analysis in Proteome Discover. B) Pie chart summarizing the proteomics dataset in terms of the impact of LPS. Orange indicates only proteins present during LPS treatment (145), yellow indicates proteins that were significantly upregulated due to LPS (91), blue indicates proteins that were decreased due to LPS (7), and black indicates proteins that were only present in control conditions (4). C) Pathway analysis of the 145 proteins only found on the LD surface of the LDs isolated from mice treated with LPS as compared to control LDs. D) Left: hierarchical clustering of proteins significantly different between control and LPS. The threshold for clustered proteins was determined by the significance between groups. Right: proteins were further grouped using K-means statistics, breaking the control and LPS-induced PLIN2 interactions into 10 protein clusters. The clustered proteins were mapped to specific metabolic pathways using Panther classification overrepresentation testing. E) The proteins directly interacting with PLIN2 under pro-inflammatory conditions. F) Co-localization of Cy3-labeled COX2 with Cy5-labeled PLIN2 (top images) and Cy3-labeled PLA_2_ with Cy5-labeled PLIN2 (bottom images) in primary mouse hepatocytes revealed yellow-to-orange areas where both probes overlapped. FRET imaging was performed as described in the Methods section. The FRET overlay was pseudo-colored to visualize regions of higher and lower FRET as shown by the inset color scale; n=12-16 cells. Calculated FRET distances are listed as inserts in the overlay image. G) Co-immunoprecipitation of COX1, COX2, and PLA_2_ with PLIN2 in LD fractions. Quantitative immunoprecipitations were performed as described in the methods; n=3-5. Figure S8A shows quantification of the immunoprecipitation. H) Homology model of PLIN2 constructed by taking the previously reported homology model based on the PLIN3 crystal structure (PDB entry 1AEP) and refining it with SWISS-Modeler. The model includes the four-helix bundle, the proposed fatty acid-binding pocket, and the 11-mer repeat/PAT domain. Based on this model, we modeled lipid binding in silico and report it in the results and Figure S8B. I-M) Tryptophan quenching binding assays were performed to validate the *in silico* lipid binding. PLIN2 binding affinity for I) arachidonic acid (AA), J) docosahexaenoic acid (DHA), K) prostaglandin E2 (PGE_2_), L) prostaglandin H2 (PGH_2_), and M) prostaglandin F2A (PGF_2a_). Binding constants (K_d_) are shown in the insets. ND= not detected.

We confirmed the immunoprecipitation proteomics by employing the APEX-biotin ligase approach described previously (Figure S7).^35,36^ In these experiments, fusion constructs expressing APEX-PLIN2, APEX-PLIN5, APEX-H2A (nuclear-targeted), and APEX-MTS (mitochondrial-targeted) were transduced into mouse embryonic fibroblasts (MEFs), treated with PBS or LPS, and the APEX-biotin ligase was triggered (Figure S7A-C). Each construct labeled between 2500-2800 proteins (Figure S7E). Previous proteomics studies of organelles in which APEX2 was exposed to the cytosol or different organelle compartments have reported non-specific labeling of cytosolic proteins.^35–37^ This limitation was addressed by using additional constructs to control for non-specific labeling at each cellular interface where PLIN2 could potentially interact. Overlaying each APEX-fusion construct indicated that PLIN2 specifically interacted with 1192 distinct proteins that PLIN5, H2A, or the mitochondria did not interact with (Figure S7F). Comparing these 1192 proteins to our PLIN2 immunoprecipitation experiment, we observed an 87% overlap. Pathway analysis mapped LPS-enriched APEX-PLIN2-specific protein IDs to pathways including cytochrome P450 metabolism, arachidonic acid metabolism, and energy/lipid metabolism, matching the immunoprecipitation results (Figure S7G-H). LPS decreased PLIN2 interactions with proteins related to metabolic processing, including protein metabolic processes (amino acid metabolism), cAMP signaling (fatty acid oxidation via lipolysis), mRNA surveillance, and ribosome biogenesis (Figure S7G). String analysis highlighted that the link between the APEX-PLIN2 and PLIN2 immunoprecipitation datasets was arachidonic acid signaling and oxylipin production (Figure S7I).

### PLIN2 directly interacts with PLA_2_ and COX2 at the LD surface

To establish PLIN2’s ability to directly interact with enzymes involved in arachidonic acid metabolism and inflammation, we performed FRET imaging with PLIN2 and COX2 using acceptor photobleaching, as described in our previous work.^38,39^ FRET was observed between PLIN2 and COX2, and the mean distance *R* between the pair was calculated to equal 65±1 Å (Figure 3F), indicating proximity. To visualize where in the cell FRET occurred, FRET efficiency (*E*) images were generated (Figure 3F; colored image). In COX2/PLIN2-labeled cells, high-intensity areas identified by morphology as LD showed E in the range of 45-60% (blue to cyan on the FRET inset color scale), consistent with efficient overlap of electronic states and direct interaction of PLIN2 with COX2 on the LD surface. FRET imaging was also performed to determine direct protein-protein interaction between PLIN2 and the arachidonic acid-promoting enzyme PLA_2_ (Figure 3F). FRET was observed between PLIN2 and PLA_2,_ and the mean distance *R* between the pair was 51±5 Å (Figure 3F). We confirmed interaction between PLIN2 and PLA_2_/COX-2 via co-precipitates with PLIN2 in pull-down experiments (Figure 3G and S8A). In response to LPS, PLIN2-COX2 and PLIN2-PLA_2_ interactions increased 2.4- and 3.8-fold, respectively, while interaction with COX1, the anti-inflammatory cyclooxygenase, decreased 4.7-fold (Figure S8A). These results indicate PLIN2 coordinates efficient pro-inflammatory oxylipin production via its interaction with PLA_2_ and COX2.

### PLIN2 binds arachidonic acid but not downstream metabolites

Given the results of the FRET and interactome experiments, we tested PLIN2’s ability to bind the product of PLA_2_, arachidonic acid, and the downstream products of COX2, prostaglandin E2 (PGE_2_), prostaglandin H2 (PGH_2_), and prostaglandin F2α (PGF2α). Using our previously published 3D homology model^38^ (Figure 3H and S8B), refined using the SWISS-MODEL AlphaFold protein-folding algorithm, we docked arachidonic acid (Figure S8B) and confirmed PLIN2 side chains in three conformational states, yielding ΔG binding of -20.4, -13.6, and -7.2 kcal/mol (Figure S8C). Predicted binding was not observed with PGH_2_, PGF_2a_, or PGE_2_. Using our published binding assay,^38,39^ we experimentally confirmed that PLIN2 bound arachidonic acid and docosahexaenoic acid with K_d_ values of 22 nM and 55 nM, respectively, while no binding was detected with the arachidonic acid metabolite PGE2, PGH2, or PGF_2a_ (Figure 3I-M). These results suggest that PLIN2 acts as a scaffold and transfer protein between PLA_2_ and COX-2 on the LD surface.

### LPS causes hyperacetylation of LD proteins

Under basal conditions, PLIN2 is not known to contribute to inflammation. Therefore, we sought to determine how PLIN2 and the LD proteome transition from benign to proinflammatory. To do so, we characterized the LD acetylome (Figure 4) and phosphoproteome (Figure 5). Using the PTMscan acetyl enrichment kit, acetylated LD proteins were enriched to 46% (Figure 4A). This yielded 4,248 acetylated proteins from 40,251 peptide spectra. LPS-induced LDs showed a significant enrichment of 120 acetylated proteins (Figure 4B), of which 63 acetylated proteins only targeted the LD during LPS treatment (Figure 4C; orange). Additionally, the number of modifications on a given protein increased under LPS conditions (Figure 4D). Pathway analysis mapped the 63 LPS-only acetylated LD proteins to generalized immune and stress responses (Figure 4E). The remaining 57 acetylated proteins that were enriched on the LD surface due to LPS mapped broadly to inflammation-related pathways (Figure 4E).

**Figure 4.**
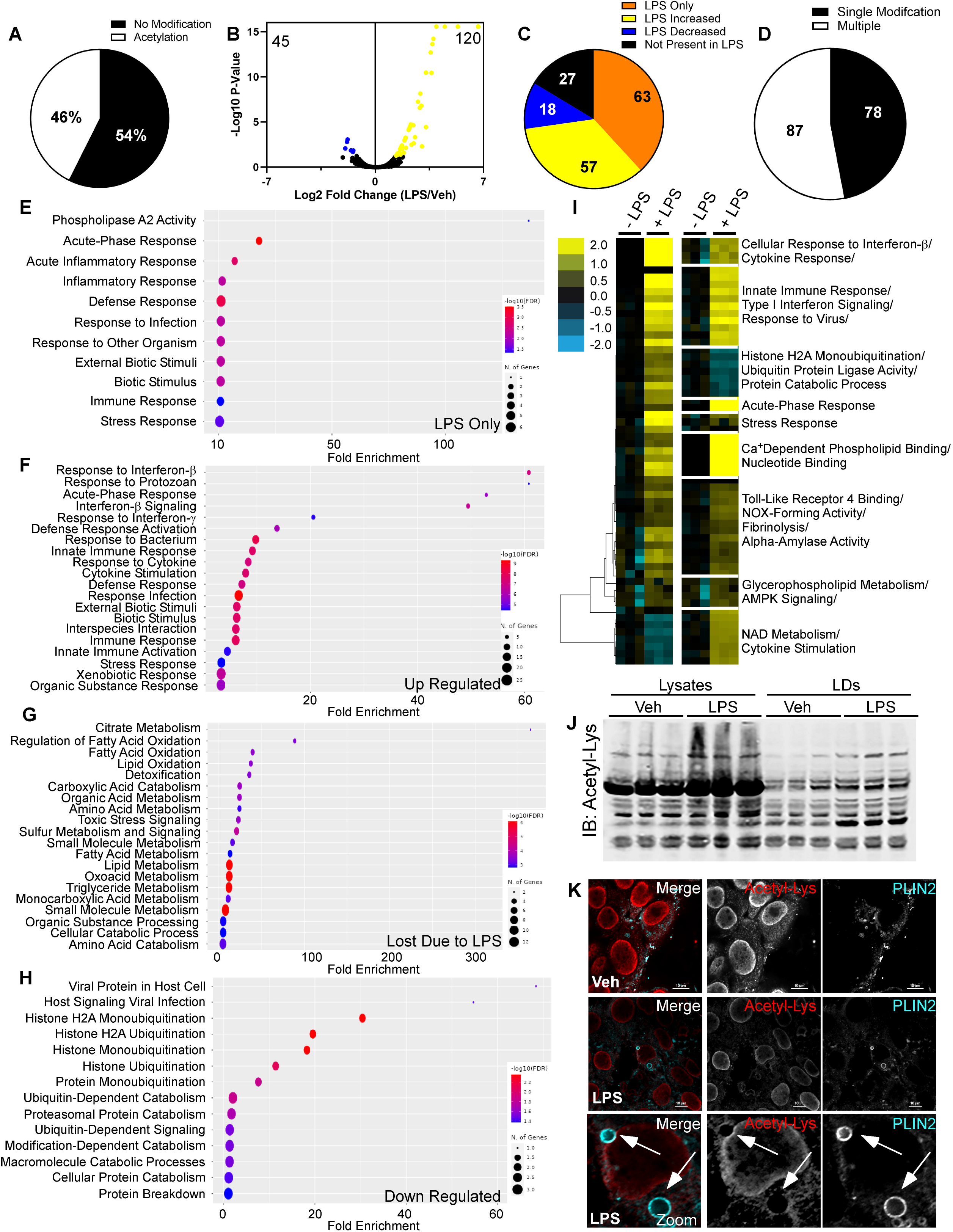
Hyperacetylation of LD proteins occurs during LPS-stimulated inflammation. A) Unlabeled quantitative acetylomics was performed on control and LPS-induced LDs. The enrichment of acetylated peptides for the proteomics dataset was 46.8% at the upper end of the expected enrichment for PTMScan-Acetylome. B) Volcano plot summarizing the changes in the acetylated peptides of LDs isolated from the livers of control and LPS-treated mice; n=3. Blue indicates proteins that were enriched in the control LD fractions, and yellow indicates LPS-enriched proteins. Significance was determined using non-parametric quantitative analysis in Proteome Discover. C) Pie chart summarizing the proteomics dataset in terms of the impact of LPS. Orange indicates targeting the LD only during LPS treatment, yellow indicates proteins that were significantly upregulated due to LPS, and blue indicates proteins that were decreased due to LPS. D) Pie chart summarizing the number of acetylation events detected in the statistically significant proteins due to LPS. E-H) Pathway analysis for the 165 statistically significant proteins in LPS-induced LDs. F) Left: hierarchical clustering of proteins significantly different between control and LPS-induced LDs replicate fractions. The threshold for clustered proteins was determined by the significance between groups. Right: proteins were further grouped using K-means clustering, dividing the significant control and LPS-induced LD proteins into 10 protein clusters. We mapped the clustered proteins to specific metabolic pathways using Panther classification overrepresentation testing. J) Hepatic lysates and LD fractions were analyzed by SDS-PAGE and immunoblotting for total acetylated lysine. Samples normalized to Ponceau S are shown in Figure S8A-B. K) Co-localization of Cy3-labeled PLIN2 with Cy5-labeled pan-acetylated lysine in ALM12 hepatocytes treated with PBS or LPS. Arrows indicate distinct changes in acetylation patterns at or near the LD surface; quantification in Figure S9C. n=4-6 images with 6-8 cells in each image.

**Figure 5.**
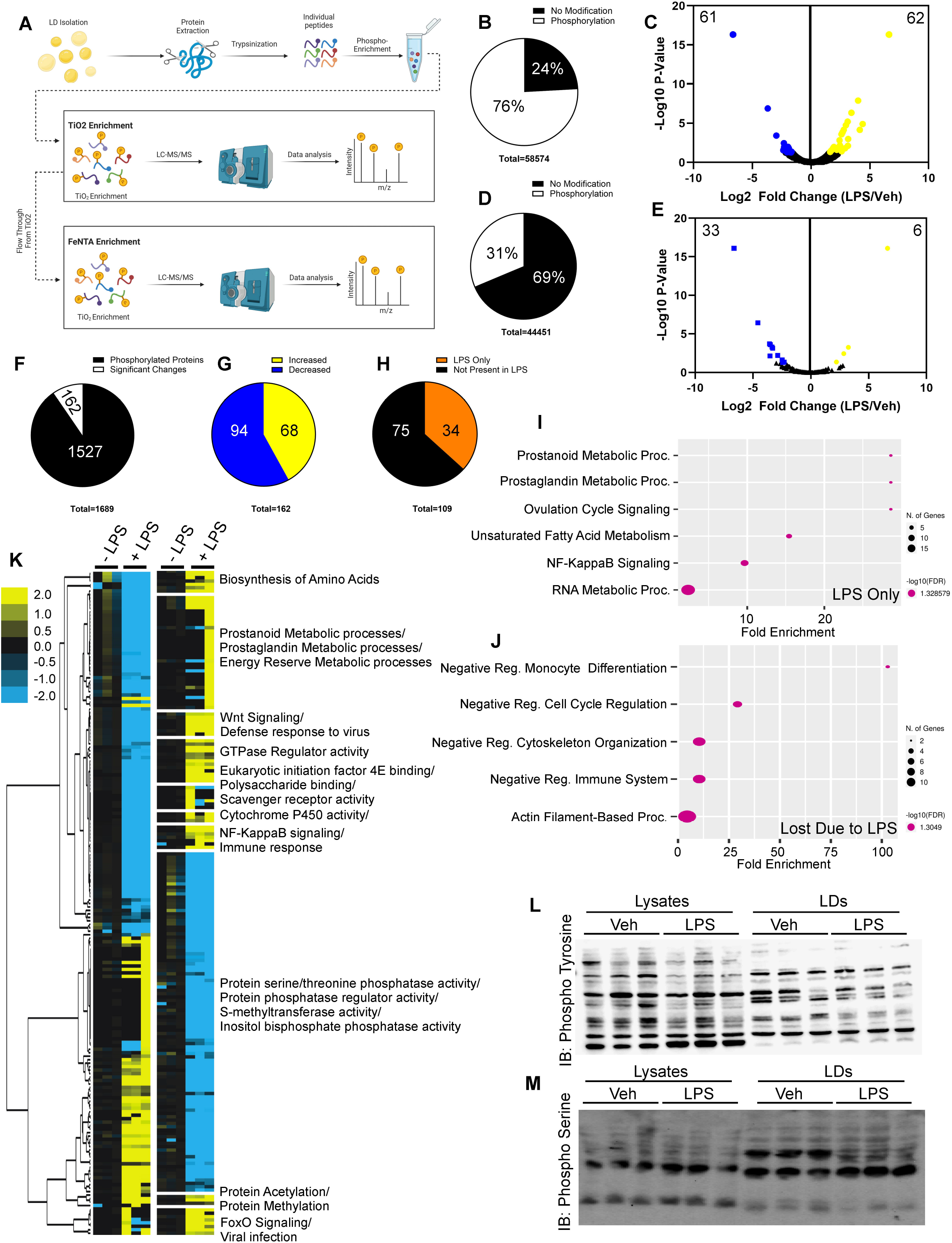
LPS-simulated inflammation suppresses the LD phospho-proteome. A) Unlabeled quantitative phosphoproteomics was performed on control and LPS-induced LDs. Schematic representation of the SMOAC workflow that was used for our proteomics approach. B) The enrichment of phosphorylated peptides for the TiO2 enrichment proteomics dataset was 76%. C) Volcano plot summarizing the changes in the TiO2-enriched peptides of LDs isolated from the livers of control and LPS-treated mice. D) Second enrichment using FeNTA, resulted in an enrichment of only 31%. E) Volcano plot summarizing the changes in the FeNTA-enriched peptides of LDs isolated according to the diagram in panel (A); n =3. Blue indicates proteins that were enriched in the control LD fractions, and yellow indicates LPS-enriched proteins. We determined significance using non-parametric quantitative analysis in Proteome Discover. F) Pie chart summarizing the impact of LPS on the proteome. Black represents the total number of phosphorylated peptides detected, while white represents the number of significantly changed phosphorylation events in the combined dataset. G-H) Pie chart summarizing the proteomics dataset in terms of statistical changes due to LPS. Yellow indicates increased LD phosphorylation due to LPS, while blue indicates a decrease. Orange indicates proteins targeted to the LD only during LPS treatment, and black indicates phosphorylated proteins not detected in LPS. I-J) Pathway analysis for the 162 statistically significant proteins. K) Left: hierarchical clustering of proteins significantly different between control and LPS-induced LD replicate fractions. The threshold for clustered proteins was determined by the significance between groups. Right: proteins were further grouped using K-means clustering, dividing the significant control and LPS-induced LD proteins into 10 clusters. We mapped the clustered proteins to specific metabolic pathways using Panther classification overrepresentation testing. L-M) Hepatic lysates and LD fractions were analyzed by SDS-PAGE and immunoblotting for phospho-tyrosine (L) and phospho-serine (M). Samples normalized to Ponceau S are shown in Figure S9E.

In contrast, proteins lost and downregulated with LPS treatment largely mapped to metabolic pathways (Figure 4G-H). Hierarchical clustering highlighted specific sets of proteins differentially enriched in control and LPS-acetylated LD proteins (Figure 4I). Using KEGG pathway analysis, statistically significant proteins that were clustered together and were increased in LPS-treated samples mapped to cellular response to interferon-γ and cytokine signaling, the innate immune response, calcium-dependent phospholipid binding and metabolism, TLR4 signaling, and acute phase response (Figure 4I; yellow). Genes that were downregulated due to LPS clustered around histone ubiquitination, protein catabolism, and glycerol lipid metabolism (Fig 4I; blue). Western blot analysis of total hepatic lysates as well as isolated LDs confirmed the hyperacetylation of proteins both in general and at the LD surface due to LPS-induced inflammation (Figure 4J and Figures S9A-B). Inflammation-induced LD hyperacetylation was strong enough to be detected by immunohistochemistry (Figure 4K and S9C-D). While acetylated proteins were observed across the cytoplasm in control conditions (Figure 4K; top panel and S8C-D), acetylated protein signal changed dramatically with LPS: the cytosolic signal localized into ring structures that colocalized with PLIN2 (Figure 4K; middle and bottom panels). Colocalization analysis indicated that in response to LPS, PLIN2 is localized with 88% of the acetylated signal around the LD (Figure S9D). These results indicate that PLIN2-LDs become hyperacetylated during inflammation, suggesting that acetylation of the LD proteome drives the LD switch from a metabolic organelle to a pro-inflammatory sink.

### LD protein phosphorylation decreases due to LPS

In contrast to the dramatic LD acetylome changes observed, LPS had minimal changes to the phospho-proteome (Figure 5). Using the SIMOC method^40–42^ for phospho-protein enrichment (Figure 5A), we enriched the proteomes to 76% and 31% via the TiO2 and FeNTA enrichment kits, respectively; TiO2 preferentially enriches phospho-tyrosine, while FeNTA enriches phospho-serine (Figure 5B and C). This yielded 1689 phosphoproteins from 103,025 peptide spectra. Of the 1689 phosphorylated proteins, only 162 were significantly changed due to LPS (Figure 5D). Across the individual enrichment steps, TiO2 enrichment resulted in 62 increased and 61 decreased (Figure 5E), while FeNTA enrichment resulted in 6 increased and 33 decreased (Figure 5F), meaning 67 were increased due to LPS, and 94 were decreased. Pathway analysis mapped the 67 LPS-increased phosphoproteins to prostanoid metabolism, prostaglandin metabolism and signaling, unsaturated fatty acid metabolism, NF-κB signaling, and RNA metabolic processing (Figure 5I).

LPS decreased 94 phospho-LD proteins, primarily associated with negative regulation of immune function, cell cycle progression, and cytoskeletal organization (Figure 5J). Hierarchical clustering revealed distinct phosphoproteomic signatures between control and LPS-treated LDs (Figure 5K; yellow). LPS-increased proteins were largely enriched in metabolic and inflammatory pathways, whereas decreased proteins were associated with immune repression and phosphatase/methyltransferase activity (Figure 5K; blue). These findings contrast with the increased protein acetylation and acyltransferase activity observed in Figure 4. Consistent with the phosphoproteomic analysis, western blotting of total hepatic lysates and isolated LDs revealed minimal changes in global or LD-associated phosphorylation following LPS-induced inflammation (Figure 5L–M and S9E–G). Thus, PLIN2-coated LDs do not undergo widespread hyperphosphorylation during inflammation. Instead, LPS selectively alters phosphorylation of phospholipid-metabolizing enzymes and NF-κB signaling components while suppressing phosphorylation associated with metabolism and negative regulation of innate immunity.

### PLIN2 is hyperacetylated under LPS conditions

Across our proteomics experiments, we identified acetylation as the only PTM changing on PLIN2 (Figure 6A). While others have reported PLIN2 phosphorylation under other metabolic conditions,^7,43^ acetylation was the only PLIN2 PTM identified under LPS conditions. PLIN2 had seven modified residues: lysine 140, 202, 243, 271, 287, 362, and 416 (Figure 6A). All seven of the residues showed increased acetylation (202, 362, 416) or became de novo acetylated with LPS (140, 243, 271, 287) (Figure 6A). PLIN2 acetylation was low in hepatic tissue under basal conditions, but LPS increased acetylation 2.6-fold (Figure 6B-C). All but one of the acetylated lysine residues reside in the C-terminal fatty acid binding pocket of PLIN2 (Figure 6D). Lysine 140, located in the N-terminal region of PLIN2, has been reported to regulate PLIN2 proteasomal degradation.^44,45^ The other six acetylation events are novel modifications. To determine which of the acetylation events were responsible for PLIN2’s proinflammatory phenotype, we generated lysine-to-arginine mutant constructs at each of the six acetylated lysine residues. We transfected these constructs into primary MEFs isolated from PLIN2 KO mice. PLIN2 KO MEFs showed blunted inflammation when exposed to LPS in culture (Figure 6E-I), while transfection of eGFP-PLIN2 restored inflammatory signaling (Figure 6E-H). Similar inflammatory signaling was observed using PLIN2 mutant constructs K140R, K202R, and K243R, indicating that these acetylation sites did not regulate PLIN2 inflammatory signaling properties. In contrast, K287R and K362R suppressed LPS-induced inflammation across most measured markers, while K271 and K416 showed variable results. Given these results, PLIN2 acetylation at the tail end of the C-terminus in the hinge region of the 4-helix bundle appears to be critical for its proinflammatory properties (Figure 6D). This region overlaps with the known PLIN5 phosphorylation site (S155), which is critical for altering PLIN5 fatty acid binding and localization.^39^ Taken together, PLIN2 K271, K287, K362, and K416 acetylation are critical residues for PLIN2 to convey inflammatory signaling.

**Figure 6.**
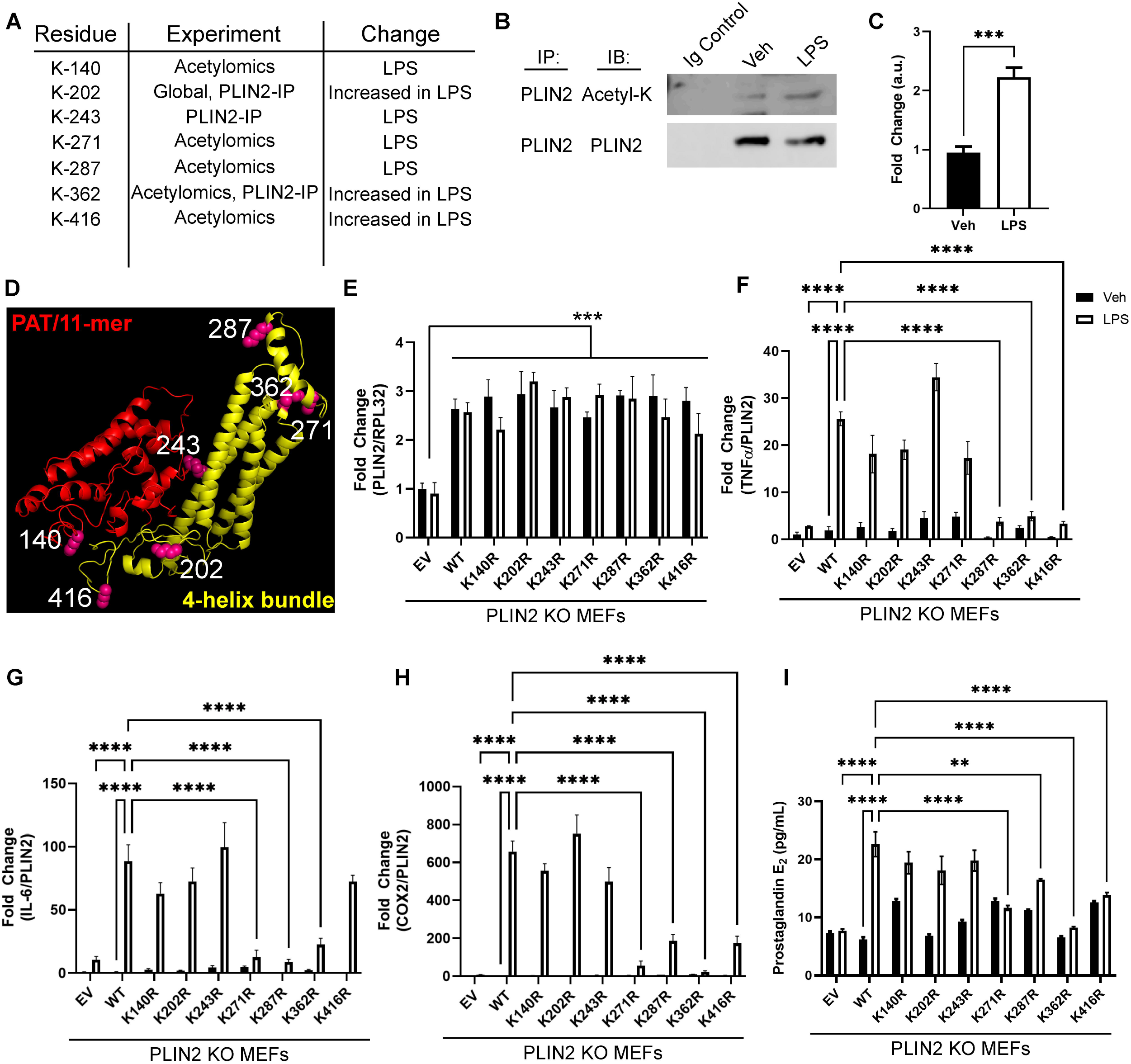
PLIN2 Acetylation is critical for proinflammatory signaling. A) Summary of acetylated residues in PLIN2 and the experimental conditions in which the modification was detected. B) Immunoprecipitation of PLIN2 from livers of control and LPS-treated mice; n = 3-4 per group. C) Quantification of PLIN2 acetylation due to LPS. ***indicates statistical significance between groups as determined by Student’s t-test. D) Homology model of PLIN2 as determined by SwissModeler and refined by Phyre3. Red indicates the PAT and 11-mer repeat region previously shown to anchor PLIN2 to the LD surface, yellow highlights the αβ-domain and 4-helix bundle responsible for PLIN2 lipid binding, and purple highlights the 7 lysines shown to be acetylated in our proteomics datasets. E) PLIN2 expression levels were determined for PLIN2 KO MEFs transfected with eGFP-PLIN2 and lysine mutants K140R, K202R, K243R, K271R, K287R, K362R, and K416R. F-H) Expression of select pro-inflammatory genes in PLIN2 KO MEFs transfected with eGFP-PLIN2 and lysine mutants: TNFα (F), IL-6 (G), and COX2 (H). Genes were normalized to PLIN2 expression levels after being normalized to the endogenous control RPL32. I) ELISA quantifying PGE_2_ in lysates of PLIN2 KO MEFs transfected with eGFP-PLIN2 and lysine mutants shown to harbor acetylation due to LPS; n=4-6 per transfection. Significance was determined by two-way ANOVA. **P<0.01, ***P<0.005, ****P<0.001.

### Acetylation of PLIN2 K271, K287, K362 and K416 drives hepatic inflammation

Based on our findings that acetylated lysine 271, 287, 362, and 416 are critical for PLIN2 proinflammatory effect, we generated eGFP-PLIN2 WT and acetylation-dead-quad mutant (QM; K271R, K287R, K362R, K416R) viruses and transduced them into PLIN2 KO mice (Figure 7A-B). Rescue of PLIN2 WT and QM resulted in detectable levels of eGFP-PLIN2 at the protein (Figure 7A) and RNA levels (Figure 7B). We also examined the acetylation state of PLIN2 WT and QM after LPS treatment (Figure 7C). Mutation of residues 271, 287, 362, and 416 resulted in a loss of detectable acetylation of the QM construct (Figure 7C). As a result of the decreased acetylation, PLIN2 KO mice transduced with QM showed a blunted response to LPS-induced inflammation (Figure 7D-H). PLIN2 KO mice rescued with the WT construct responded to LPS, demonstrated by induction of inflammatory cytokines (Figure 7D), PLA_2_ activity, and PGE_2_ levels (Figure 7E-F). In the PLIN2 KO mice rescued with the eGFP-QM construct, LPS treatment did not induce inflammatory gene expression, cPLA_2_ activity, or PGE_2_ production (Figure 7D-F). Protein abundance of ALOX12, 5-LOX, and COX2 was significantly increased in LD fractions of eGFP-WT-expressing mice treated with LPS (Figure 7G-H), unlike the QM mice (Figure 7G-H). These data indicate that acetylation of PLIN2 at residues 271, 287, 362, and 416 is responsible not only for the expression of innate immune and oxylipin proteins, but also for the targeting of inflammatory proteins to the LD surface to propagate the inflammatory response.

**Figure 7.**
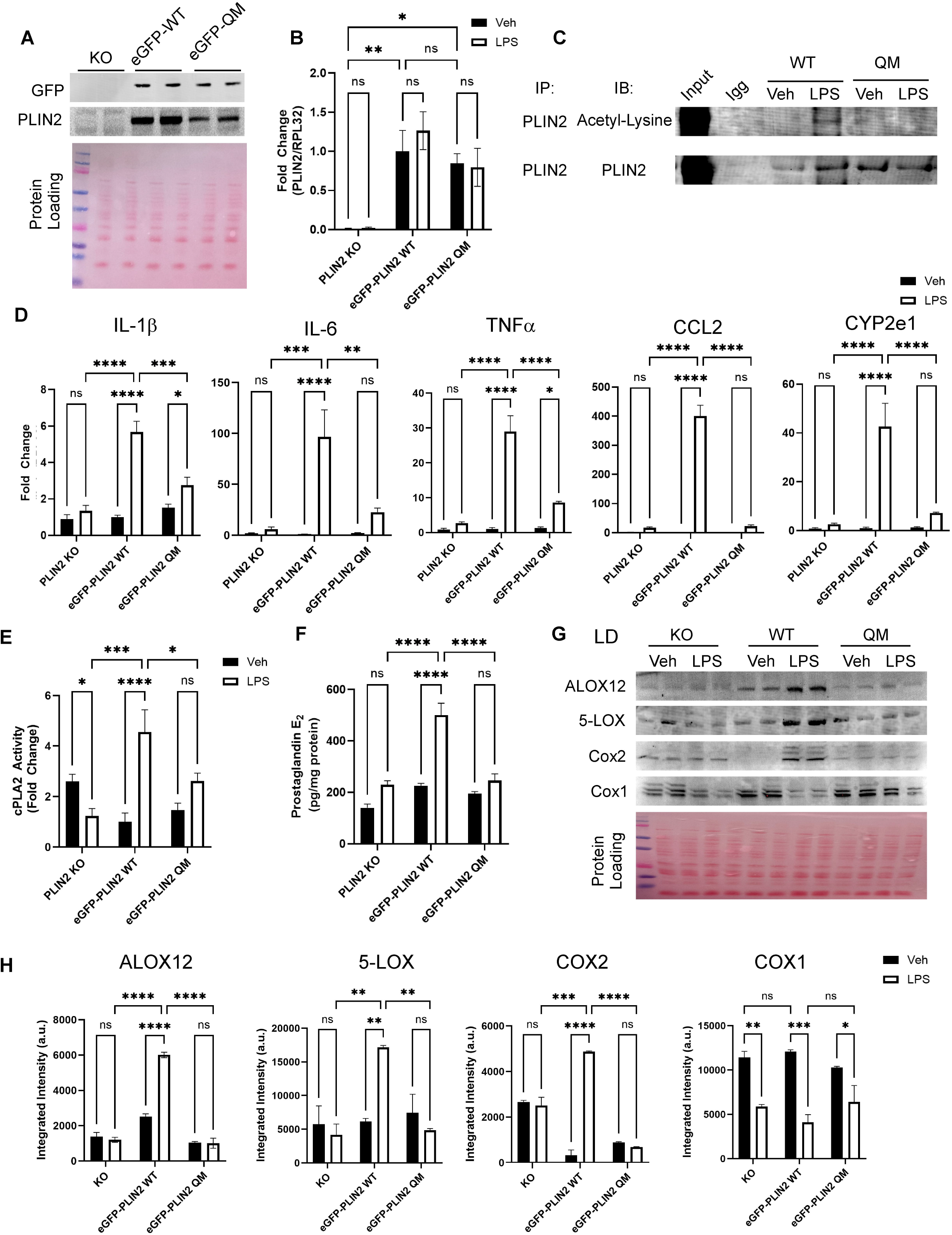
PLIN2 acetylation dead mutants block LPS-induced inflammation *in vivo*. A) PLIN2 KO mice were transduced with eGFP-PLIN2 or eGFP-PLIN2 QM (K271R, K287R, K362R, and K416R) AAVs. Expression of PLIN2 was determined via Western blotting for PLIN2 and eGFP. B) RNA levels from livers of PLIN2 KO mice transduced with eGFP-PLIN2 or eGFP-QM were determined. C) The acetylation state of WT and QM PLIN2 due to LPS was determined via immunoprecipitation. D) Expression of select pro-inflammatory genes was determined in the livers of PLIN2 KO, PLIN2 KO-eGFP-WT, and PLIN2 KO-eGFP-QM mice treated with or without LPS; n=4. *P<0.05, **P<0.02, ***P<0.01, ****P<0.001. E) cPLA_2_ activity was determined from livers and normalized to eGFP-PLIN2 WT-expressing mice treated with PBS; n=4. *P<0.05, **P<0.02, ***P<0.01. ****P<0.001. F) PGE_2_ in hepatic lysates of PLIN2 KO, PLIN2 KO-eGFP-WT, and PLIN2 KO-eGFP-QM mice treated with or without LPS; n=4. Significance was determined by two-way ANOVA. LDs isolated from PLIN2 KO, PLIN2 KO-eGFP-WT, and PLIN2 KO-eGFP-QM mice treated with or without LPS were subjected to Western blot analysis. LDs represent two pooled fractions, with 4 mice per treatment. **P<0.01, ***P<0.005, ****P<0.001.

## DISCUSSION

The above data show that inflammation promotes PLIN2 expression and LD accumulation, which in turn recruits oxylipin-synthetic enzymes involved in the innate immune response to the LD surface. This recruitment drives a sustained inflammatory phenotype in which cytokines and arachidonic acid-derived oxylipins are produced at high levels. Hyperacetylation of the LD proteome drives recruitment, triggering a cascade of proinflammatory signals that match several disease states, including Alzheimer’s, Prion Disease, NASH/MASH, and viral/bacterial infection. Our results indicate PLIN2 coordinates part of the inflammatory phenotype. The transfer of arachidonic acid from cPLA_2_ to oxylipin-producing enzymes such as COX2, 5-LOX, and ALOX-12 appears to be the primary mechanism of action for PLIN2. This model is supported by PLIN2’s ability to directly interact with cPLA_2_ and COX2 and bind AA, but not prostaglandins. Removing PLIN2 or acetylating it blocks this step in the signaling cascade. Ablation of PLIN2 or its acetylation also suppresses the induction of the innate immune response, suggesting additional mechanisms of action that remain to be investigated.

It has been proposed that membrane-bound organelles, such as LDs, recruit and assemble key adaptor molecules to support the innate immune response and influence the intensity and speed of pathway activation.^1,46^ For example, LDs act as anchors for critical signaling steps mediating TLR-Viperin induction of IFN-γ.^46–48^ LDs have also been shown to recruit key innate immune proteins that act as antimicrobial/antiviral sinks.^1^ Thus, LDs are increasingly recognized to act as a major regulatory checkpoint in the innate immune system. Our findings indicate that PLIN2 has a mechanistic role in the innate immune signaling cascade. Ablation of PLIN2 suppresses critical cytokine and innate immune signaling proteins associated with NF-κB activation. Given the link between arachidonic acid-based oxylipin products and innate immune signaling,^31,49^ PLIN2 likely plays a critical role in coupling oxylipin production to NF-κB signaling.

Conversely, PLIN2 overexpression leads to LD accumulation in numerous tissues, including the heart, liver, and muscle.^7^ While many mechanisms likely underlie the different effects of PLIN2 metabolic regulation, its role in inflammation is undoubtedly a major contributor. PLIN2 expression and the coinciding accumulation of LDs are increased in response to LPS or other inflammatory stimuli. PLIN2 expression is also increased in tissues or cells characterized by chronic inflammation,^50^ including microglia from aged mice,^18,51^ foam cells and atherosclerotic plaques,^7,52^ non-alcoholic steatohepatitis livers,^12,50^ cancers,^8,9^ and glial cells of Alzheimer’s patients.^21,23,53–55^ PLIN2 ablation attenuates, and overexpression exacerbates LPS-mediated inflammation.^10,12,24^ Tissue-specific or global PLIN2 deletion broadly reduces inflammation associated with obesogenic diets and protects against atherosclerosis.^10,12,50^ While correlated with inflammation, the underlying mechanism linking PLIN2 to inflammation has largely been unexplored. Our findings not only confirm that PLIN2 ablation protects against inflammation but also detail a mechanism in which acetylated PLIN2 interacts with the oxylipin enzymes PLA2 and COX2, as well as the innate immune system at the LD surface, propagating an inflammatory cascade. Further linking PLIN2 and inflammation, recombinant PLIN2 binds various lipids but shows the highest affinity for arachidonic acid. These results suggest that PLIN2 acts as a scaffold and transfer protein between PLA_2_ and COX-2 on the LD surface. Because LDs have emerged as major sites of oxylipin generation and are commonly correlated with inflammation, these data implicate PLIN2 as a key player in this lipid-mediated signaling and the innate immune response.

## ACKNOWLEDGEMENTS

We thank the University of Minnesota Imaging Center, the Center for Metabolomics and Proteomics, the University of Pittsburgh Health Sciences Mass Spectrometry Core, and the Virginia Tech metabolism core for providing instrumentation and technical support. We thank Joshua Drake, Ann Hertzel, Todd Lydic, David Berhnlor, Siobhan Craige, Stella Volpe, Eva Schmelz, and Barbara Atshaves for scientific discussions. Funding was provided for Charles Najt (NIH: R00AG070104, Pratt Foundation), Mahima Devarajan (NIH: F30AG076322), Alex Churchill (Virginia Tech IGEP: Translational Obesity and Health Span Fellowship), Samuel Gerrard (USDA: GRANT1398196), Stacy Gelhaus (NIH: S10OD023402, S10OD032141), and Douglas Mashek (NIH: R01DK132849 and R01DK114401). The following instrumentation grant was critical for the development of new proteomics methodologies as part of the University of Minnesota Center for Metabolomics and Proteomics (NIH: S10OD02717).

## AUTHOR CONTRIBUTIONS

CPN, DGM conception and design of research; CPN, MD, EK, AC, LB, JR, and SG performed experiments; CPN, MD, CG, LAH, JR, AC, and SG analyzed data; CPN, SG, and DGM interpreted results of experiments; CPN prepared figures; CPN drafted the manuscript; CPN, SG, JH, KM, LH, and CG contributed materials and reagents necessary for the completion of studies; CPN, DGM, MD, LAJ, DG, and EK edited and revised the manuscript; All authors approved the final version of the manuscript.

## DECLARATION OF GENERATIVE AI AND AI-ASSISTED TECHNOLOGIES

The authors declare that they did not use generative AI or AI-assisted technologies in the submitted manuscript.

## STAR METHODS AND KEY RESOURCES

### Key Resource Table

| REAGENT or RESOURCE | SOURCE | IDENTIFIER |
| --- | --- | --- |
| **Antibodies** |  |  |
| Rabbit polyclonal anti-PLIN2 | (Atshaves BP et al., 1999) | N/A |
| Mouse polyclonal anti-PLIN2 | Protein Tech | 60340-1-Ig |
| Guinea pig polyclonal anti-PLIN3 | Progen | GP37 |
| Guinea pig polyclonal anti-PLIN5 | Progen | GP31 |
| Mouse polyclonal anti-COX1 | Cayman Chemical | 160109 |
| Mouse polyclonal anti-COX2 | Cayman Chemical | 160126 |
| Mouse polyclonal anti-PGES | Cayman Chemical | 160140 |
| Rabbit polyclonal anti-cPLA<sub>2</sub> | Cell Signaling Technologies | 2832 |
| Rabbit polyclonal anti-P-cPLA<sub>2</sub> | Cell Signaling Technologies | 2831 |
| Rabbit polyclonal anti-COXIV | Cell Signaling Technologies | 4850 |
| Rabbit monoclonal anti-H3 | Cell Signaling Technologies | 4499 |
| Mouse monoclonal anti-AcLysine | Novis Biologicals | 15G10 |
| Mouse monoclonal anti-AcLysine | Cell Signaling Technologies | 9681 |
| Mouse monoclonal anti-AcLysine | Santa Cruz | AKL5C1 |
| Mouse monoclonal anti-AcLysine | ThermoFisher | 1C6 |
| Mouse monoclonal anti-phospho serine | ThermoFisher | MA1-91608 |
| Mouse monoclonal anti-phospho tyrosine | MilliporeSigma | P1869 |
| Mouse monoclonal anti-PLIN2 | ProteinTech | 6034Ig |
| Rabbit monoclonal anti-5-lipoxygenase | Cell Signaling Technologies | 3289 |
| Rabbit polyclonal anti-5-lipoxygenase | Protein Tech | 10021-1-ig |
| Rabbit monoclonal anti-ALOX12 | Cell Signaling Technologies | 49874 |
| Mouse monoclonal anti-GFP | Cell Signaling Technologies | 2955 |
| Mouse monoclonal anti-alpha-tubulin | Cell Signaling Technologies | 3873 |
| Mouse monoclonal anti-PGE<sub>2</sub> | Cayman Chemical | 414013 |
| Rabbit monoclonal anti-Phospho-STAT3 | Cell Signaling Technologies | 9145 |
| Mouse monoclonal anti-Total-STAT3 | Cell Signaling Technologies | 9139 |
| Rabbit monoclonal anti-Viperin | Cell Signaling Technologies | 75654 |
| Rabbit polyclonal anti-IGTP | Cell Signaling Technologies | 14979 |
| Donkey anti-Guinea pig IRDye 800CW | LI-COR | Cat No. 926-32411 |
| Donkey anti-Guinea pig IRDye 680RD | LI-COR | Cat No. 926-68030 |
| Donkey anti-Rabbit pig IRDye 800CW | LI-COR | Cat No. 926-32213 |
| Donkey anti-Rabbit pig IRDye 680RD | LI-COR | Cat No. 926-68022 |
| Donkey anti-Mouse pig IRDye 800CW | LI-COR | Cat No. 925-32212 |
| Donkey anti-Mouse pig IRDye 680RD | LI-COR | Cat No. 926-68023 |
| **Bacterial and Virus Strains** |  |  |
| DH5a | Invitrogen | 18258-012 |
| XL1-Blue | Agilent | Cat No. 200150 |
| Stable | NEB | C3040H |
| AAV8SP[CMV]-eGFP-PLIN2 | VectorBuilder |  |
| AAV8SP[CMV]-eGFP-PLIN2-QM | VectorBuilder |  |
| Lenti-CMV-APEX2-PLIN2 | This paper |  |
| Lenti-CMV-APEX2-PLIN5 | This paper |  |
| Lenti-CMV-APEX2-MTS | This paper |  |
| Lenti-CMV-APEX2-H2A | This paper |  |
| M15 | Najt et al 2014 |  |
| **Chemicals, peptides, and recombinant proteins** |  |  |
| Trypsin Gold, Mass Spec Grade | Promega | V5280 |
| Dynabeads Protein G | ThermoFisherScientific | 10004D |
| NucBlue | ThermoFisherScientific | R37605 |
| LPS (Stereotype: O111:B4) | MilliporeSigma | L2630 |
| BioTracker CoxFluor | MilliporeSigma | SCT079 |
| Red/Green BODIPY PC-A2 | ThermoFisherScientific | A10072 |
| Biotin-xx-tyramide | MilliporeSigma | SML3484 |
| Ponceau S | Fisher BioReagents | BP103-10 |
| AutoDOT-Neutral lipid Stain | Abcepta | SM1000a |
| hPLIN2 Recombinant proteins | Najt CP, et al., 2014 |  |
| **Critical Commercial Assays** |  |  |
| Nuclear Complex Co-IP Kit | Active Motif | 54001 |
| Proteome Profiler XL Cytokine Array | R&D Systems | ARY028 |
| PTMScan Acetyl-Lysine Motif Kit | Cell Signaling Technologies | 13416 |
| Effectene | QIAGEN | Cat No. 301425 |
| High-Select TiO<sub>2</sub> | ThermoFisherScientific | A32993 |
| High-Select Fe-NTA | ThermoFisherScientific | A32992 |
| QuickChange Lightning Site-Directed | Agilent | 210518 |
| **Deposited Data** |  |  |
| LD Global Proteomics | In progress at Massive |  |
| LD Global Acetylomics | In progress at Massive |  |
| PLIN2-IP Proteomics | In progress at Massive |  |
| LD Global Phospho-proteomics | In progress at Massive |  |
| PLIN2-APEX Interactome | In progress at Massive |  |
| **Experimental Models: Cell Lines** |  |  |
| AML12 | ATCC | CRL-2254 |
| *Plin2* Null Hepatocytes | This paper |  |
| Primary Hepatocytes | This paper |  |
| *Plin2* Null MEFs | This paper |  |
| **Experimental Models: Organisms/Strains** |  |  |
| Male C57BL6/J mice | Envigo | N/A |
| *Plin2* Floxed Mice | Andrew S. Greenberg<sup>50</sup> | N/A |
| *Plin2* Global Knockout mice | This Paper | N/A |
| B6.FVB-Tg(EIIa-Cre)C5379Lmgd/J Cre mice | Jax Laboratory |  |
| **Recombinant DNA** |  |  |
| EX-Mm20284-M29-eGFP-PLIN2 | GeneCopoeia |  |
| eGFP-PLIN2 K140R | This paper |  |
| eGFP-PLIN2 K202R | This paper |  |
| eGFP-PLIN2 K243R | This paper |  |
| eGFP-PLIN2 K271R | This paper |  |
| eGFP-PLIN2 K362R | This paper |  |
| eGFP-PLIN2 K416R | This paper |  |
| pLenti-CMV-APEX2-PLIN2 | This paper |  |
| pLenti-CMV-APEX2-PLIN5 | This paper |  |
| pLenti-CMV-APEX2-MTS | This paper |  |
| pLenti-CMV-APEX2-H2A | This paper |  |
| pAAV8SP[CMV]-eGFP-PLIN2 | VectorBuilder |  |
| pAAV8SP[CMV]-eGFP-PLIN2-QM | VectorBuilder |  |
| pQE9-His-PLIN2 | (Najt CP et al., 2014) |  |
| **Software and Algorithms** |  |  |
| SigmaPlot 11 | Systat Software, Inc | N/A |
| Prism8 | GraphPad | N/A |
| Canvas 11 | Canvas | N/A |
| Avizo 3D Software | Thermo Fisher Scientific | N/A |
| Image Studio v5 | LI-COR Biosciences | N/A |
| NIS-Elements 4 | Nikon | N/A |
| Microsoft Photos Application | Microsoft | N/A |

## METHODS

## LEAD CONTACT AND MATERIALS AVAILABILITY

The lead contact for this study is Charles Najt. Studies were performed in collaboration and with partial support from Douglas Mashek. All unique/stable reagents generated in this study are available from either Drs. Najt or Mashek with a complete Materials Transfer Agreement.

### Mice

The University of Minnesota and Virginia Tech Institutional Animal Care and Use Committee approved all animal protocols. Male 8-12-week-old C57BL6/J mice were obtained from Harlan Laboratories, while *Plin2* floxed mice were a kind gift from Andrew S. Greenberg. ^50^ All mice were housed under controlled temperature and lighting (20-22°C; 14:10-h light-dark cycle). The mice were fed a purified control diet (TD 94045; Harlan Teklad Premier Laboratory) and acclimatized for 1 week before any experimental procedure. PLIN2 global knockout mice were generated by crossing *Plin2* floxed mice with B6.FVB-Tg(EIIa-Cre)C5379Lmgd/J Cre mice obtained from Jax Laboratories. Ablation of *Plin2* was determined via PCR and Western blotting (Fig 4A). For acute inflammation studies, C57BL6/J, *Plin2* floxed, and *Plin2* knockout mice were injected with sterile saline or 5 mg/kg LPS (Stereotype: O111:B4) overnight. Before harvest, mice were fasted for 4 h. All mice were euthanized using ketamine/xylazine for liver tissue and serum collection.

### Antibodies

The following antibodies were used in this study: rabbit polyclonal anti-PLIN2 (1:3000 dilution; (Atshaves BP et al., 1999), 1:1000 dilution; ProteinTech 60340-1-1g), guinea pig polyclonal anti-PLIN3 (1:1000 dilution; Progen GP37), guinea pig polyclonal anti-PLIN5 (1:1000 dilution; Progen GP31), mouse polyclonal anti-COX1(1:1000 dilution; Cayman Chemical 160109), mouse polyclonal anti-COX2 (1:1000 dilution; Cayman Chemical 160126), mouse polyclonal anti-PGES (1:1000 dilution; Cayman Chemical 160140), rabbit polyclonal anti-PLA2 (1:500 dilution; Cell Signaling Technologies 2832), rabbit polyclonal anti- and phospho-PLA2 (Ser505) (1:500 dilution; Cell Signaling Technologies 2831), rabbit polyclonal anti-COXIV (1:1000 dilution; Cell Signaling Technologies 4850), rabbit monoclonal anti-H3 (1:1000 dilution; Cell Signaling Technologies 4499), mouse monoclonal anti-AcLysine (1:500 dilution; Novis Biologicals 15G10), mouse monoclonal anti-AcLysine (1:500 dilution, Santa Cruz Biotechnology AKL5C1), mouse monoclonal anti-AcLysine (1:1000 dilution, Cell Signaling Technology 9681S), mouse monoclonal anti-AcLysine (1:1000 dilution, ThermoFisher 1C6), mouse monoclonal anti-phosphoserine (1:500 dilution, ThermoFisher MA1-91608), mouse monoclonal anti-phosphotyrosine (1:500 dilution, MilliporeSigma P1869), mouse monoclonal anti-biotin (1:1000 dilution, MilliporeSigma SAB4200680), donkey anti-Guinea pig IRDye 800CW (1:10,000 dilution, LI-COR 926-32411), donkey anti-guinea pig IRDye 680RD (1:10,000 dilution, LI-COR 926-68030), donkey anti-rabbit IRDye 800CW (1:10,000 dilution, LI-COR 926-32213), donkey anti-rabbit IRDye 680RD (1:10,000 dilution, LI-COR 926-68022), donkey anti-mouse IRDye 800CW (1:10,000 dilution, LI-COR 925-32212), donkey anti-mouse IRDye 680RD (1:10,000 dilution, LI-COR 926-68023), donkey anti-mouse secondary antibody HRP (1:30,000 dilution, ThermoFisher A16011).

### Plasmids

For overexpression or rescue experiments the following plasmids were purchased or generated: EX-Mm20284-M29-eGFP-PLIN2 (purchased from GeneCopoeia), EX-Mm20284-M29-eGFP-PLIN2 K140R (generated by site-directed mutagenesis using QuickChange Lightning Site-Directed Mutagenesis Kit, Agilent), EX-Mm20284-M29-eGFP-PLIN2 K202R (generated by site-directed mutagenesis using QuickChange Lightning Site-Directed Mutagenesis Kit, Agilent), EX-Mm20284-M29-eGFP-PLIN2 K243R (generated by site-directed mutagenesis using QuickChange Lightning Site-Directed Mutagenesis Kit, Agilent), EX-Mm20284-M29-eGFP-PLIN2 K271R (generated by site-directed mutagenesis using QuickChange Lightning Site-Directed Mutagenesis Kit, Agilent), EX-Mm20284-M29-eGFP-PLIN2 K287R (generated by site-directed mutagenesis using QuickChange Lightning Site-Directed Mutagenesis Kit, Agilent), EX-Mm20284-M29-eGFP-PLIN2 K362R (generated by site-directed mutagenesis using QuickChange Lightning Site-Directed Mutagenesis Kit, Agilent), EX-Mm20284-M29-eGFP-PLIN2 K416R (generated by site-directed mutagenesis using QuickChange Lightning Site-Directed Mutagenesis Kit, Agilent), pLenti-CMV-APEX2-PLIN2 (pLenti-CMV backbone was obtained from AddGene 17448, pcDNA FlagAPEX2 NES was a gift from Alice Ting. A fusion of APEX2-PLIN2 was PCR amplified and cloned into the pLenti-CMV backbone), pLenti-CMV-APEX2-PLIN5 (PLIN5 was PCR amplified out of the mCherry-PLIN5 plasmid from ^39^ and subcloned into pLenti-CMV-APEX2 in place of PLIN2), pLenti-CMV-APEX2-MTS (A fusion of APEX2-MTS was PCR amplified using the following MTS sequence described in ^56^), pLenti-CMV-APEX2-H2A (A fusion of APEX2-H2A was PCR amplified and cloned into the pLenti-CMV backbone. The H2A construct was subcloned from the sequence described in ^56^).

### LD Isolation

All procedures were performed using pre-chilled equipment and solutions. The following method was adapted from ^57,58^. Livers from C57BL6/J at 12-16 wk of age were injected with saline or LPS [5mg/kg] were harvested, rinsed in PBS, minced, and suspended in 6 ml LD-lysis buffer [10 mM Tris-base pH 7.0 with protease (Complete protease inhibitor cocktail, Roche, Basel Switzerland), phosphatase (PhosSTOP, Sigma-Aldrich, St. Louis, MO), and deacetylase inhibitors (De-acetylase Cocktail, MedChem Express, Monmouth Junction, NJ)]. The resuspended hepatic tissue was mechanically disrupted with 15 strokes in a glass Teflon Dounce homogenizer. The homogenate was then transferred to a 15 ml Falcon tube and centrifuged in an Avanti J-15R (Beckman Coulter) swinging bucket rotor at 900xg for 10 min at 4°C. The post-nuclear supernatant (PNS), including the fat cake, was loaded onto a sucrose step gradient (4 ml 35% sucrose in LD-Lysis buffer, 4 ml 15% sucrose in LD-lysis buffer, 4 ml PNS) and centrifuged at 96,000xg in a Beckman SW41ti swinging bucket rotor for 4 h. The LD fraction appeared as a white film at the top of the tube, which was removed with a Pasteur pipette. LDs were snap-frozen or processed immediately. Additional cellular fractions were collected and analyzed for fraction purity and protein movement via western blotting.

### Quantitative global proteomics

Samples for quantitative proteomic analysis were collected from four control and four LPS-treated mice. LDs from these mice were harvested as described above. The LDs appeared as a white lens at the top of the tube. Samples were collected using a Pasture pipette. For proteomic analysis, LD samples were analyzed using the following protocol with minor modifications.^59^ Proteins were extracted with proteomic lysis buffer [PLB1; 7 M urea, 2 M thiourea, 0.4 M tris pH 8, 20% acetonitrile, 10 mM tris (2-carboxyethyl) phosphine (TCEP), 40 mM chloroacetamide]. The samples were pipetted up and down several times and vortexed. 100 μg of total LD protein from each sample determined by Bradford assay was diluted fivefold with water, and then trypsin digested at a 1:40 ratio of trypsin (Promega, Madison, WI) to total protein. Samples were incubated overnight for 16 h at 37°C. After incubation, the samples were frozen at – 80°C and dried in a vacuum concentrator. Each sample was cleaned with a 1 CC Waters Oasis MCX cartridge (Waters Corporation, Milford, MA), and the eluate was dried *in vacuo*. Samples were resuspended in water, and peptide concentration was determined by BCA assay. Equal peptide amounts for each sample were desalted on a C18 stage tip ^60^ and reconstituted for analysis.

We reconstituted the dried peptide fractions in 97.9:2:0.1, H2O:acetonitrile (ACN):formic acid (FA) (load solvent) and analyzed ∼600 nanograms by capillary LC-MS with a Thermo Fisher Scientific, Inc (Waltham, MA) Dionex UltiMate 3000 RSLCnano system on-line with an Orbitrap Eclipse mass spectrometer (Thermo Scientific, Waltham MA) with FAIMS (high-field asymmetric waveform ion mobility) separation. We injected peptides directly in load solvent and performed gradient separation on a self-packed C18 column (Dr. Maisch GmbH ReproSil-PUR 1.9 um 120 Å C18aq, 100 um ID x 40 cm length) at 55 °C with the following profile: 5% B solvent from 0 – 2 minutes, 8% B at 2.5 minutes, 32% B at 100 minutes, 45% B at 120 minutes and 90% B at 125 minutes with a flowrate of 425 nl/min from 0 – 2 minutes and 325 nl/minute from 2.5 – 125 minutes, where solvent A was 0.1% formic acid in water and solvent B was 0.1% formic acid in ACN. The FAIMS nitrogen cooling gas setting was 5.0 L/min, the carrier gas was 4.6 L/min and the inner and outer electrodes were set to 100°C. We scanned the CV (compensation voltage) at -50 and -70 for 1 second each with a data-dependent acquisition method. We employed the following MS parameters: ESI voltage +2.1 kV, ion transfer tube 275 °C; no internal calibration; Orbitrap MS1 scan 120k resolution in profile mode from 380 – 1400 *m/z* with 50 msec injection time; 100% (4 x 10^5^) automatic gain control (AGC); high energy collisional dissociation (HCD) MS2 activation was triggered on precursors with 2 – 5 charges above 1x10^4^ counts; MIPS (monoisotopic peak determination) was set to Peptide; MS2 settings (both CV’s) were: 1.6 Da quadrupole isolation window, 30% fixed collision energy, ion trap detection with rapid scan mode, 35 msec max injection time, 100% (1 x 10^4^) AGC, 45 sec dynamic exclusion duration with +/- 10 ppm mass tolerance and exclusion lists were shared among CV’s.

We processed peptide tandem MS using SEQUEST (Thermo Scientific) in Proteome Discoverer 2.5. The mouse Universal Proteome (UP000000589) database was downloaded from UniProt.org on 11/19/19 and merged a common lab contaminant protein database (ftp://ftp.thegpm.org/fasta/cRAP) (55,180 total protein sequences). We applied the precursor mass recalibration node with precursor mass tolerance 20 ppm, product ion tolerance 0.6 Da and fixed carbamidomethyl (CAM) modification of cysteine 57.0215 *m/z*. The SEQUEST (PMID: 24226387) database search parameters were: enzyme trypsin full specificity, 2 missed cleave sites; precursor tolerance 15 ppm, fragment ion tolerance was 0.6 Da. We specified CAM cysteine (+57.021 Da) as a fixed modification and the dynamic modifications were oxidation of methionine (+15.995 Da), acetylation of lysine (+42.011 Da), phosphorylation of serine, threonine and tyrosine (+79.966 Da), conversion of glutamine to pyroglutamic acid (-17.027 Da), deamidation of asparagine or glutamine (+0.984 Da), acetylation of protein N-terminus (+42.011 Da), methionine loss at the protein N-terminus (-131.040 Da) and methionine loss + acetylation at the protein N-terminus (-89.030 Da). We used the Percolator algorithm with a concatenated target-decoy database approach to control the false discovery rate. (https://doi.org/10.1038/nmeth1113) We reported protein lists with a 1% FDR threshold.

For each file, we applied the untargeted Minora Feature Detector algorithm in PD 2.4 for the assignment of chromatographic features for isotopically related peaks within a 0.2-minute retention time range. After feature assignment, we performed chromatographic alignment; we used the file with the most features as the reference and aligned features from each file to it with a 5-minute (RT) window tolerance and 12 ppm mass tolerance. Retention times were adjusted using a non-linear regression-based fit of the differences in retention time between the reference and the sample. Peptides were mapped to retention time-aligned consensus features across samples with the requirement that at least one sample contains a peptide spectral match.

For each sample, protein abundances were calculated by summing the abundances of consensus features for related peptides. All peptide forms were used for protein quantification calculations. We applied normalization across samples using total peptide abundances. For each sample, protein abundances were summed and adjusted with a normalization factor, which is the quotient of sample abundance divided by the abundance of the reference sample, defined as the sample in the experiment with the maximum abundance. The normalization factors were used for adjustment of peptide and protein group abundances for the entire protein report. We set the hypothesis test method to t-test and we reported p-values that were adjusted using the Benjamini-Hochberg correction for false discovery rate.^61^ Controlling the false discovery rate: a practical and powerful approach to multiple hypothesis testing.

### Heatmap and pathway analysis of proteomics data

Pathway analysis, hierarchical clustering, and heat map production were carried out as previously described ^12,62–67^. The hierarchical clustering and heat maps were generated using Cluster 2.0 from the Eisen Laboratory, modified by Michiel de Hoon (http://bonsai.hgc.jp/~mdehoon/software/cluster/). Java Tree Viewer was used to view and color the heat map.

### Western blotting

Cell lysates (30-80 µg protein) were separated on 10-12% tricine gels using a Mini-Protean II cell (Bio-Rad, Hercules, CA) system at constant amperage (30 mA per gel) for about 4 h. Proteins were then transferred onto PVDF membranes at constant voltage (90 V) for 2 h. Blots were stained with Ponceau S to confirm uniform protein loading ^68,69^ before blocking in 5% BSA or 7% non-fat milk in TBST (10 mM Tris-HCl, pH 8, 100 mM NaCl, 0.05% Tween-20) for 1 h. Blots were incubated with specific poly- or monoclonal antibodies overnight and were developed with IRDye 800CW (LI-COR) or IRDye 680RD (LI-COR) secondary antibodies. To visualize the bands of interest, blots were scanned using the LI-COR Odyssey imaging system (Lincoln, NE). Protein bands were quantified by densitometric analysis after image acquisition using NIH Scion Image to obtain relative protein levels expressed as integrated density. All values were normalized to Ponceau S staining ^70–73^.

### Serum analysis

Serum β-hydroxybutyrate measurements were determined from mouse serum samples using a β-hydroxybutyrate LiquiColor kit (Stanbio Laboratory, Boerne, TX, USA) according to the manufacturer’s instructions. Liver TG levels were determined using Wako Chemicals lipid assay systems (Wako Diagnostics, Richmond, VA). Colorimetric analyses of lipids were measured at 570nm on an Omega FLOUstar 96-well plate reader from BMG Labtech (Ortenberg, Germany).

### Fatty acid oxidation using [^14^C]-oleate

Methods for measuring FA oxidation were adapted from Huynh et al.^74^ and Najt et al.^59^ In brief, liver lysates were suspended in mitochondrial respiration buffer (MRB; 100 mM Sucrose, 10 mM Tris-HCL pH 7.4, 5 mM potassium phosphate, 0.2 mM EDTA, pH 8.0, 80 mM potassium chloride, 1 mM magnesium chloride, 2 mM carnitine, 0.1 mM malate, 0.05 mM coenzyme A, 2 mM ATP, 1 mM DTTpH8.0). Samples [100 μg] (50 μL suspension) were added to 350 µL MRB supplemented with 0.7% FA-free BSA/500 μM oleate/0.5μCi [^14^C]-oleate. Reactions were incubated at 37 °C for 1 h, quenched by adding 200 µL of 1 M perchloric acid, vortexed, and then centrifuged at >14,000 × g for 15 min at room temperature. 400 μL of the supernatant was transferred to a scintillation vial, and the radioactive acid-soluble products were counted on a scintillation counter.

### Cell culture

All cells were maintained in a humidified incubator at 37°C under 5% CO_2_. We obtained AML-12 cells from ATCC (Manassas, VA). Cells were grown in DMEM: F12 (1:1) supplemented with 10% FBS, ITS (10 mg/ml insulin, 5.5 mg/ml transferrin, 5 ng/ml selenium), and 40 ng/ml dexamethasone. Primary hepatocytes were isolated as described previously.^39,75^ Primary hepatocytes were cultured at 37°C under 5% CO_2_ in M199 medium containing 23 mM HEPES, 26 mM sodium bicarbonate, 10% FBS, 50 IU/ml penicillin, 50 μg/ml streptomycin, 100 nM dexamethasone, 100 nM insulin, 11 mM glucose, and 1 mM carnitine [Base M199 Media]. We isolated primary mouse embryonic fibroblasts (MEFs) with minor adaptations (^76^). MEFs were harvested at embryonic day 13.5-14.5 and, upon plating, were maintained in hypoxic conditions for three serial passages before being placed under 5% CO_2_ for experimental conditions. Base media for the primary MEF cultures consisted of DMEM/F10 (1:1) supplemented with nonessential amino acids, 10% FBS, and Glutamax [MEF Media]. For imaging experiments, cells were treated under one of the following conditions: control media (Base M199 Media, MEF Media, AML12 Media), or LPS media (Base M199 Media, MEF Media, or AML12 Media supplemented with 1 μg/ml LPS). For LD counterstaining experiments, control and LPS media were supplemented with AutoDOT neutral lipid stain (Absepta) 20 min before imaging. For COX2 activity experiments, cells were treated with LPS media overnight. Before imaging, cells were treated with CoxFluor dye for 1 h.^25^ In a subset of the CoxFluor assays, cells were transfected with eGFP-PLIN2 and select mutants. For experiments with BODPIY PC-A2 phospholipase A2 substrate, 30 μl of 10 mM DOPC (1,2-dioleoyl-snglycero-3-phosphocholine; Avanit Polar lipids), 30 μl of 10 mM DOPG (1,2-dioleoyl-sn-glyero-3-[phospho-rac-(1-glycerol)]; Avanit Polar Lipids), and 30 μl of 1 mM PC-A2 (each prepared using ethanol as the solvent) were mixed. 77 ml of this ethanolic lipid mixture was pipetted rapidly into 5 ml of 50 mM Tris, 100 mM NaCl, 1 mM CaCl_2_ (pH 8.8) with vortexing for 2 min.^30^ This liposomal incorporated substrate mixture was added to cells 6 h prior to treatment with PBS or LPS media.

### Live cell imaging

We performed fluorescence imaging experiments with cells seeded at 50,000 cells/plate on Mat-Tek cover-glass plates (Ashland, MA). Primary hepatocytes, MEFs, and AML12 cells were treated with the corresponding media, counterstained for LDs, and/or treated with the COX2 activity sensor, CoxFluor, or the PLA2 sensor, PC-A2. Digital images were acquired using a Nikon A1Rsi Laser Scanning Confocal Imaging System (LSCIS; Nikon, Melville, NY) equipped with 405 nm, 488 nm, 561 nm, and 640 nm laser, four-channel GaSP detectors, and a 63x water immersion objective (University of Minnesota Imaging Center). Images of the cells were acquired by sequential excitation. Image files were analyzed using NIS-Elements software or FIJI-Image software. Z-stack or multiple focal planes were imaged to ensure compartmentalization and localization.

### EicosaCell Assay

The EicosaCell assay was performed following the procedure of Bandeira-Melo et al. ^26^ In brief, LPS-treated primary mouse hepatocytes were permeabilized and fixed in 1% EDAC for 30 min at 37°C. The cells were washed with HBSS and blocked with 2% FBS in HBSS for 1 hour before incubation with primary antibodies [mouse anti-PGE_2_ at 1:100 dilution and rabbit anti-Plin2 at 1:50 dilution] for 1 h at room temperature. After extensive washing with HBSS, Cy3 or Cy5 labeled secondary antibodies (at 1:100 dilutions) in PBS were added to each set and incubated for 1 h at room temperature. Cells were then washed with HBSS and mounted with cover slips using fluorogel mounting medium (Electron Microscopy Science, Hatfield, PA). Cells were stained with Cy3 and Cy5 alone in the absence of primary antibodies to serve as background controls. LPS-treated cells fixed in 4% paraformaldehyde and permeabilized with 0.1% saponin served as negative controls for EDAC-mediated entrapment of PGE_2_ at sites of synthesis. Cell images were sequentially acquired on a Nikon A1Rsi Laser Scanning Confocal Imaging System (LSCIS; Nikon, Melville, NY) equipped with 405 nm, 488 nm, 561 nm, and 640 nm laser, four-channel GaSP detectors, and a 63x water immersion objective. Co-localization of the Cy3- and Cy-5 signals were obtained using the Nikon Elements software where the confocal images from the green and red channels were merged and appeared yellow where superimposition occurred (red and green are additive and yield yellow to orange in RGB color space).

### PLIN2 Immuno-Precipitation (IP) for quantitative mass spectrometry

LDs were isolated from saline- and LPS-treated mice as described above. A Western blot of the LD fractions was performed to determine the fold increase of PLIN2 due to LPS. The co-immunoprecipitation input was normalized to PLIN2, not total protein.^34^ This method quantifies changes in interacting proteins due to biological conditions, not due to changes in PLIN2 abundance. Prior to co-immunoprecipitation for mass spectrometry analysis, the anti-PLIN2 antibody^77^, was coupled to Dyna Protein G beads using the Dynabeads Antibody Coupling Kit (ThermoFisher). For IP from cell fractions, we incubated a minimum of 2 mg of LD proteins with 10μg of indicated primary antibody coupled Dyna beads in 750 μL IP buffer (20mM HEPES pH 7.4, 0.15 M NaCl, 1 mM EDTA,1 mM EGTA, 1 mM DTT, 1x protease inhibitor cocktail, 1x phosphatase inhibitor cocktail, and 1x deacetylase inhibitor cocktail) for 16 h at 4°C with tumbling. The beads were washed three times with cold IP buffer. Input samples were kept for further analysis. IP samples were eluted with 100 μl 0.2 M glycine (pH 2.6) and incubated at room temperature for 10 min. A second elution was performed, and the two eluents were combined, neutralized with an equal volume of 1 M Tris pH 8.5, and snap-frozen.

IP samples were thawed, and protein amount was determined by Bradford assay. 50 μg of protein was reduced and alkylated with 5 mM DTT and 15mM iodoacetamide in the dark for 1 h. The reaction was then diluted fivefold with 50 mM Tris pH 8, then trypsin digested at a 1:40 ratio of trypsin (Promega, Madison, WI) to total protein. Samples were incubated overnight (16 h) at 37°C. After incubation, the samples were frozen at -80°C and dried in a vacuum concentrator. Each sample was desalted on a C18 stage tip ^60^ and reconstituted for analysis. We reconstituted the dried peptide fractions in 97.9:2:0.1, H2O:acetonitrile (ACN):formic acid (FA) (load solvent) and analyzed ∼600 nanograms by capillary LC-MS with a Thermo Fisher Scientific, Inc. (Waltham, MA) Dionex UltiMate 3000 RSLCnano system on-line with an Orbitrap Eclipse mass spectrometer (Thermo Scientific, Waltham, MA) with FAIMS (high-field asymmetric waveform ion mobility) separation. We injected peptides directly in load solvent and performed gradient separation on a self-packed C18 column (Dr. Maisch GmbH ReproSil-PUR 1.9 um 120 Å C18aq, 100 um ID x 40 cm length) at 55 °C with the following profile: 5% B solvent from 0 – 2 minutes, 8% B at 2.5 minutes, 32% B at 100 minutes, 45% B at 120 minutes and 90% B at 125 minutes with a flowrate of 425 nl/min from 0 – 2 minutes and 325 nl/minute from 2.5 – 125 minutes, where solvent A was 0.1% formic acid in water and solvent B was 0.1% formic acid in ACN. The FAIMS nitrogen cooling gas setting was 5.0 L/min, the carrier gas was 4.6 L/min and the inner and outer electrodes were set to 100°C. We scanned the CV (compensation voltage) at -50 and -70 for 1 second each with a data-dependent acquisition method. We employed the following MS parameters: ESI voltage +2.1 kV, ion transfer tube 275°C; no internal calibration; Orbitrap MS1 scan 120 k resolution in profile mode from 380 – 1400 *m/z* with 50 msec injection time; 100% (4 x 10^5^) automatic gain control (AGC); high energy collisional dissociation (HCD) MS2 activation was triggered on precursors with 2 – 5 charges above 1x10^4^ counts; MIPS (monoisotopic peak determination) was set to Peptide; MS2 settings (both CV’s) were: 1.6 Da quadrupole isolation window, 30% fixed collision energy, ion trap detection with rapid scan mode, 35 msec max injection time, 100% (1 x 10^4^) AGC, 45 sec dynamic exclusion duration with +/- 10 ppm mass tolerance and exclusion lists were shared among CV’s.

We processed peptide tandem MS using SEQUEST (Thermo Scientific) in Proteome Discoverer 2.5. The mouse Universal Proteome (UP000000589) database was downloaded from UniProt.org on 11/19/19 and merged with a common lab contaminant protein database (ftp://ftp.thegpm.org/fasta/cRAP) (55,180 total protein sequences). We applied the precursor mass recalibration node with precursor mass tolerance 20 ppm, product ion tolerance 0.6 Da and fixed carbamidomethyl (CAM) modification of cysteine 57.0215 *m/z*. The SEQUEST (PMID: 24226387) database search parameters were: enzyme trypsin full specificity, 2 missed cleave sites; precursor tolerance 15 ppm, fragment ion tolerance was 0.6 Da. We specified CAM cysteine (+57.021 Da) as a fixed modification and the dynamic modifications were oxidation of methionine (+15.995 Da), acetylation of lysine (+42.011 Da), phosphorylation of serine, threonine and tyrosine (+79.966 Da), conversion of glutamine to pyroglutamic acid (-17.027 Da), deamidation of asparagine or glutamine (+0.984 Da), acetylation of protein N-terminus (+42.011 Da), methionine loss at the protein N-terminus (-131.040 Da) and methionine loss + acetylation at the protein N-terminus (-89.030 Da). We used the Percolator algorithm with a concatenated target-decoy database approach to control the false discovery rate. (https://doi.org/10.1038/nmeth1113) We reported protein lists with a 1% FDR threshold.

For each file, we applied the untargeted Minora Feature Detector algorithm in PD 2.4 to assign chromatographic features to isotopically related peaks within a 0.2-minute retention time range. After feature assignment, we performed chromatographic alignment; we used the file with the most features as the reference, aligning features from each file with a 5-minute (RT) window tolerance and 12 ppm mass tolerance. Retention times were adjusted using a non-linear regression-based fit of the differences in retention time between the reference and the sample. Peptides were mapped to retention time-aligned consensus features across samples with the requirement that at least one sample contains a peptide spectral match.

For each sample, protein abundances were calculated by summing the abundances of consensus features for related peptides. All peptide forms were used for protein quantification calculations. We applied normalization across samples using total peptide abundances. For each sample, we summed protein abundances and adjusted them using a normalization factor: the ratio of the sample abundance to the reference sample abundance, defined as the sample with the maximum abundance in the experiment. The normalization factors were used to adjust peptide and protein group abundances for the entire protein report. We set the hypothesis test method to t-test and we reported p-values that were adjusted using the Benjamini-Hochberg correction for false discovery rate.^78^ Controlling the false discovery rate: a practical and powerful approach to multiple hypothesis testing.

### Fluorescence resonance energy transfer (FRET) imaging

Colocalization and FRET imaging studies were performed following procedures described earlier ^79^. Briefly, mouse primary hepatocytes were fixed in cold acetone/ethanol (70:30 v/v) and washed with PBS. The cells were blocked with 2% BSA and then incubated with either mouse anti-PLA_2_ (1:10) or mouse anti-COX2 (1:25) and then rabbit anti-Plin2 (1:50)] for 1 h at room temperature. After extensive washing with PBST (0.05% Tween100 in PBS), a mixture of secondary reagents consisting of Cy3- or Cy5-labeled secondary antibodies (1:100 dilution) in PBS was added to each set and incubated for 1 h at room temperature. Cells were then washed with PBS and mounted with cover slips using fluorogel mounting medium. Cells stained with Cy3 and Cy5 alone in the absence of primary antibodies served as background controls. Cell images were sequentially acquired and analyzed on an Olympus FluoView 1000 Laser Scanning Confocal Microscope as described in the previous section. Acceptor photobleaching FRET experiments were performed to measure the increase in donor (Cy3) emission upon photobleaching of the acceptor (Cy5) as described elsewhere ^79,80^. To calculate the FRET efficiency (*E*), representing the efficiency of energy transfer between donor and acceptor, the following equation was used: *E* = 1-(*I*_DA_/*I*_D_) where *I*_DA_ is the donor fluorescence intensity before acceptor photobleaching and *I*_D_ is the donor fluorescence intensity after acceptor photobleaching. An average *E* value was calculated from the increase in Cy3 fluorescence emission after photobleaching. The intermolecular distance *R* between Plin2 and COX2 or Plin2 and PLA_2_ was calculated from the equation *E* = 1/(1-(*R*/*R*_o_)^6^), where *E* is experimentally determined and *R*_o_ is the Foster radius for the Cy3-Cy5 FRET pair, previously determined as 51 Å. For the FRET efficiency images, analysis was performed in MetaMorph 7.5 (Molecular Devices, Sunnyvale, CA). Images were filtered to remove randomized noise by using a low pass filter. The filtered images of the donor emission before acceptor photobleaching were subtracted from the image after acceptor photobleaching. The resultant image was divided by the image of donor emission after acceptor photobleaching and multiplied by 100 to generate bar-scale FRET efficiencies.

### Co-immunoprecipitation studies

The Nuclear Complex Co-IP system from Active Motif (Carlsbad, CA) was used for co-immunoprecipitation (co-IP) experiments following the manufacturer’s protocol. Briefly, cell lysates were incubated overnight with kit reagents and antibodies (anti-PLIN2, anti-COX2, anti-COX1 or anti-PLA_2_) at 4°C with shaking. The next day, unbound fractions were separated by magnet, followed by washing and elution of the bound complex. Eluate proteins were analyzed by Western blotting. A parallel co-IP with the lysates using anti-rabbit IgG was performed to assess nonspecific binding (negative control). Input material was run in parallel to determine equal loading of the sample. Immunoprecipitations were performed to determine protein-protein interactions as well as protein modifications as described previously in ^81^.

### RNA isolation and RT-PCR analysis

RNA was extracted with Trizol from liver tissues, followed by reverse transcription with SuperScript® VILO™ cDNA Synthesis Kit (Invitrogen) to generate cDNA. Gene expression was quantified as described previously ^12,81,82^.

### Cytokine Proteome Profile

The cytokine array was performed with the Proteome Profiles mouse XL cytokine Array Kit (R&D Systems). Briefly, hepatic lysates from four wild-type, four PLIN2 KO, four wild-type, LPS-treated, and four PLIN2 KO, LPS-treated mice were combined for a 100 μg total lysate per treatment group (25 μg per sample x 4 samples =100 μg). The pooled samples were incubated with the array overnight at 4°C with rocking. Membranes were washed three times with 1X wash Buffer for 10 min each. The array was then incubated with the antibody cocktail for 1 h at room temperature. Membranes were washed three times before adding the provided chemiluminescent detection reagents (Figure S3A). Pixel intensity was quantified with ImageJ and normalized to internal standards in accordance with the manufacturer’s instructions.

### Quantitative oxylipin measurements

Hepatic tissue samples were extracted for oxylipins using a modified Bligh-Dyer method. Tissue (25-50 mg) was homogenized in MeOH containing 500 ng/ml of oxylipin internal standards, followed by centrifugation. Cleared supernatant was transferred to a glass tube to which 10% formic acid, dichloromethane (DCM, 0.5 ml), and water (0.5 ml) was added. After centrifugation, the organic phase was transferred to a clean glass tube, and a second DCM extraction was performed on the sample. This phase was combined with the first extraction, and the DCM was dried under N_2_. Saponification was performed at 90°C for 20 min with 1.25 M NaOH. The sample was extracted with 10% formic acid and DCM, the organic fraction was transferred to a clean glass vial, dried under a stream of N_2_, and stored at -80°C until analysis. Samples were reconstituted in 100 µl of MeOH for analysis. Oxylipins were analyzed by targeted liquid chromatography mass spectrometry using a Vanquish Horizon UPLC coupled to a Thermo fisher Quantiva triple quadrupole mass spectrometer (Thermo Fisher Scientific). Oxylipins were separated using a Luna C8(2) column (Phenomenex, 3 µm, 100 Å, 150 x 2 mm) over a gradient consisting of solvent A: H_2_O + 0.1% acetic acid and solvent B: ACN + 0.1% acetic acid. The selected reaction monitoring (SRM) scan mode was selected for oxylipins over the course of the gradient. The method included a table of parent and product ion masses for SRM that was segmented based over the course of the gradient to optimize the MS duty cycle. ThermoFisher XCalibur software was used for automatic and manual integration of oxylipin analytes and corresponding stable isotope internal standards. Data is reported as the peak area ratio of the analyte to the internal standard.

### Quantitative Acetyl-omics of LDs

Sample preparation for control and LPS induced LDs were carried out as described above and in accordance with our published works.^58,59^ LDs were allocated in 11 mg fractions as determined by the BCA assay. Proteins were reduced with 4.5 mM dithioerythritol (DTT) for 30 minutes at 37 °C, cooled to room temperature, alkylated with 10 mM iodoacetamide for 30 minutes in the dark, and then enzymatic protein hydrolysis was initiated by the addition of trypsin (1:40, w:w, enzyme: substrate) and allowed to proceed for 16 h at 37 °C with tumbling. The following day, trypsin was quenched by adding formic acid to bring the pH <2.0. Samples were dried down, then desalted using Waters’ HLB 5CC cartridges. Peptides were eluted and dried completely in a SpeedVac and stored at -80°C until further processed. Dried peptides were resuspended in 1.4 ml IAP buffer (50 mM MOPS/NAOH, pH 7.2, 10 mM sodium phosphate monobasic, 50 mM NaCl). Peptide amounts were determined via BCA and 10 mg peptides for each sample were taken forward for acetyl-peptide enrichment using PTMScan Acetyl-Lysine Motif Kit (Cell Signaling; 13416). In brief, 10 mg of peptide solution was added to the washed antibody-bead slurry and incubated at 4°C with tumbling for 2 h. Samples were centrifuged at 2,000xg for 60 sec and the supernatant was transferred into a clean Eppendorf tube for quality control purposes. Antibody-beads were gently resuspended in 1mL IAP via inversion, then centrifuged again. The wash step was repeated twice with IAP followed by three wash steps with HPLC-grade water. During the last water wash, the antibody-beads were spun a second time to ensure any residual wash was removed. Acetylated peptides were eluted from the beads with 0.15% TFA at room temperature. A second elution was performed, and the two elution fractions were combined. Acetylated peptides were dried down via a SpeedVac and cleaned up on a C18 stage tip.

We reconstituted the dried peptide fractions in 97.9:2:0.1, H_2_O:acetonitrile (ACN):formic acid (FA) (load solvent) and analyzed ∼400 nanograms by capillary LC-MS with a Thermo Fisher Scientific, Inc. (Waltham, MA) Dionex UltiMate 3000 RSLCnano system online with an Orbitrap Fusion mass spectrometer (Thermo Scientific, Waltham MA). We injected peptides directly in load solvent and performed gradient separation on a self-packed C18 column (Dr. Maisch GmbH ReproSil-PUR 1.9 um 120 Å C18aq, 100 um ID x 40 cm length) at 55°C with the following profile: 5% B solvent from 0 – 2 minutes, 12% B at 5 minutes, 40% B at 150 minutes, 50% B at 180 minutes and 90% B at 182 minutes with a flowrate of 400 nl/min from 0 – 2 minutes and 315 nl/minute from 2.5 – 180 minutes, where solvent A was 0.1% formic acid in water and solvent B was 0.1% formic acid in ACN. We employed the following MS parameters: ESI voltage +2.1 kV, ion transfer tube 275°C; no internal calibration; Orbitrap MS1 scan 120k resolution (at 200 *m/z*) in profile mode from 380 – 1500 *m/z* with 50 msec injection time; 100% (4 x 10^5^) automatic gain control (AGC); high energy collisional dissociation (HCD) MS2 activation was triggered on precursors with 2 – 5 charges above 1x10^E4 counts; MIPS (monoisotopic peak determination) was set to Peptide; MS2 settings were: 1.6 Da quadrupole isolation window, 30% fixed collision energy, ion trap detection, 35 msec max injection time, 100% (1 x 10^4^) AGC, 45 sec dynamic exclusion duration with +/- 10 ppm mass tolerance.

We processed peptide tandem MS using SEQUEST (Thermo Scientific) in Proteome Discoverer 2.5. The mouse Universal Proteome (UP000000589) database was downloaded from UniProt.org on 11/19/19 and merged with a common lab contaminant protein database (ftp://ftp.thegpm.org/fasta/cRAP) (55,180 total protein sequences). We applied the precursor mass recalibration node with precursor mass tolerance 20 ppm, product ion tolerance 0.6 Da and fixed carbamidomethyl (CAM) modification of cysteine 57.0215 *m/z*. The SEQUEST (PMID: 24226387) database search parameters were: enzyme trypsin full specificity, 2 missed cleave sites; minimum peptide length 6 residues, precursor tolerance 15 ppm, fragment ion tolerance was 0.6 Da. We specified CAM cysteine (+57.021 Da) as a fixed modification and the dynamic modifications were oxidation of methionine (+15.995 Da), acetylation of lysine (+42.011 Da), phosphorylation of serine, threonine and tyrosine (+79.966 Da), conversion of glutamine to pyroglutamic acid (-17.027 Da), deamidation of asparagine or glutamine (+0.984 Da), acetylation of protein N-terminus (+42.011 Da), methionine loss at the protein N-terminus (- 131.040 Da) and methionine loss + acetylation at the protein N-terminus (-89.030 Da).

### Phosphoproteomics of LDs

To enrich phosphopeptides from control and LPS-induced LDs, we isolated LDs as described above. We characterized LD phosphopeptides using the SMOAC method.^41^ For the SMOAC method, we subjected 1 mg of tryptic-digested LD proteins to High-Select TiO_2_ phosphopeptide enrichment according to the manufacturer’s protocol (ThermoFisher; A32993). The TiO_2_ eluent was saved for MS analysis, while the TiO_2_ flow-through wash fractions were pooled and enriched using the High-Select Fe-NTA enrichment kit (ThermoFisher; 32992). Both TiO_2_ and Fe-NTA eluents were cleaned up on a C18 stage tip and redissolved in 0.1% FA prior to LC-MS analysis. LC-MS analysis was carried out as described above in the Acetyl-omics of LDs section.

### Immunohistochemistry of hepatocytes

Briefly, mouse primary hepatocytes were fixed in cold acetone/ethanol (70:30 v/v) and washed with PBS. The cells were blocked with 5% BSA and then incubated with either mouse anti-acetyl lysine (1:25) or rabbit anti-Plin2 (1:50) for 1 h at room temperature. After extensive washing with PBST (0.05% Tween100 in PBS), add a mixture of Cy3- or Cy5-labeled secondary antibodies (1:100 dilution) in PBS to each set and incubate for 1 h at room temperature. Cells were then washed with PBS and mounted with cover slips using fluorogel mounting medium. Cells stained with Cy3 and Cy5 alone in the absence of primary antibodies served as background controls. Cell images were sequentially acquired and analyzed on a Nikon A1Rsi Laser Scanning Confocal Imaging System (LSCIS; Nikon, Melville, NY) equipped with 405 nm, 488 nm, 561 nm, and 640 nm lasers, four-channel GaSP detectors, and a 63x water immersion objective.

### PLIN2 structural prediction and analysis

Similar to methods used to predict the lipid-binding domain of PLIN2 ^38^ The PLIN2 structure was modeled by homology to the structure of residues 209-431 in PLIN3 from Protein Data Bank entry 1SZ1 by using Modeler, as implemented in the ModWeb Web server (https://modbase.compbio.ucsf.edu/modweb/) and the online web server Phyre^2^ (www.sbg.bio.ic.ac.uk/phyre2/).

### Site-directed mutagenesis

To generate eGFP-*Plin*2 K140R, K202R, K243R, K271R, K287R, K362R, K416R, K271R/K287R/K362R/K416R mutants, QuickChange Lightning Site-Directed Mutagenesis Kit (Agilent; Santa Clara, CA) was used in accordance with the manufacturer’s protocol.

### PGE_2_ Elisa

Mouse liver homogenates were prepared as described above. PGE_2_ concentration in hepatic lysates was measured using the PGE_2_ Elisa Kit (Cayman Chemicals) according to the manufacturer’s instructions. Samples were measured in triplicate.

### PLIN2 purification

Recombinant PLIN2 was purified as previously described with some modifications.^38,83^ Briefly, proteins were overexpressed in *E. coli* host strain M15 and grown at 37°C in 1 liter cultures containing 2X-YT medium with 100 µg/ml ampicillin until OD_600_ = 0.8, followed by induction with IPTG (1 mM). After 3 h, cells were harvested by centrifugation (4°C for 30 min at 3500 × g). To purify PLIN2, pelleted cells were resuspended in 50 mM Na_2_HPO_4_ (pH 8.0), 300 mM NaCl, and protease inhibitors, then sonicated. Lysates were centrifuged at 4 °C for 15 min at 25,000 × g, and the pellet was solubilized in 20 mM Tris-HCl (pH 7.9), 100 mM Na_2_HPO_4_, 500 mM NaCl, 20 mM imidazole, and 6 M guanidine hydrochloride and applied to a Ni-NTA column equilibrated in the same buffer. The column was sequentially washed with 20 mM Tris-HCl (pH 7.9), 100 mM Na_2_HPO_4_, 500 mM NaCl, 20 mM imidazole, 6 M urea, 10 mM Tris-HCl (pH 7.5), and 150 mM NaCl. The protein was eluted with 10 mM Tris-HCl (pH 7.5), 150 mM NaCl, and 50 mM EDTA, then checked for purity by SDS-PAGE.

### Intrinsic tryptophan fluorescence binding studies

The binding of fatty acids to PLIN2 was determined by measuring fluorescence quenching of PLIN2 tryptophan residues after addition of the ligand, as described previously ^38^. In brief, the intrinsic tryptophan fluorescence of PLIN2 (150 nM in 10 mM NaH_2_PO_4_, pH 7.5) was monitored from 300 to 400 nm after excitation at 295 nm (to minimize interference from tyrosine fluorescence) both before and after addition of increasing increments of arachidonic acid, docosahexaenoic acid, PGE_2_, PGH_2_, PGF_2a_, (fatty acid/prostaglandin were dissolved in 200 proof spectroscopically clear ethanol) using a Cary Eclipse fluorescence spectrophotometer. Data were corrected for background scatter originating from the buffer and ligand without protein present. The intrinsic tryptophan fluorescence in the presence of different concentrations of ligand was plotted as the maximum fluorescence difference (ΔF = F₀-F) vs. ligand concentration to yield a saturation curve, where F and F₀ were the measured fluorescence emission intensity of the protein solution in the presence and absence of ligand, respectively. The dissociation constant K_d_ was determined from the double reciprocal plot of the saturation curve. Linear regression of 1/[1-(F/F_max_)] versus [ligand]/(F/F_max_) yielded a slope = 1/ K_D_ and ordinate intercept = nE₀/ K_d,_ where F represented fluorescence intensity at a given concentration of ligand, F_max_ was the maximal fluorescence. E₀ was the protein concentration, and n equaled the number of ligand binding sites.

### PLIN2-APEX proteomics

For each APEX2 cell line, five 10-cm plates of cells (1,000,000 cells/plate) were treated with PBS or LPS media described above. Cells were subsequently treated with 500 μM biotin tyramide for 30 minutes. Before harvesting, we catalyzed protein biotinylation by adding 1 μM H_2_O_2_ for 60 seconds. We quenched the reaction by gently washing cells with PBS containing 10 mM sodium ascorbate and 5 mM Trolox. We harvested cells by scraping in PBS, centrifuged them, and snap-froze them.

Cell pellets were thawed and lysed in lysis buffer (100 mM Tris pH 7.4, 0.1% Triton-X, complete mini protease inhibitor cocktail, PhoSTOP phosphatase inhibitor, and deacetylase inhibitors [MedChem Express]). Cells were then sonicated and centrifuged at 4000 x g for 10 min, and the supernatant was transferred to a fresh tube. Protein concentrations were determined by the Pierce 660 assay. For biotinylated protein enrichment, 1 ul of streptavidin beads was used per 10 μg of protein. Streptavidin beads were equilibrated with 2 washes of PBS, 1 wash with 1 M KCl, 1 wash with 2 M urea and 50 mM Tris, pH 8, and 2 washes with lysis buffer prior to loading cell lysates. Add streptavidin beads to lysates and incubate overnight at 4°C with tumbling. Streptavidin beads were collected, and lysates were collected for further analysis. The streptavidin beads were then washed twice with lysis buffer, once with 1 M KCl, and twice with 2 M urea, 50 mM Tris, pH 8. For protein elution, the streptavidin beads were resuspended in 1 M urea, 1mM DTT, 0.4 μg Trypsin Gold, 50 mM Tris, pH 8. Elution was allowed to proceed for 1 h at 37°C. The beads were collected, and the supernatant was removed and placed in a fresh tube. The beads were then washed twice with 2 M urea, 50 mM Tris pH 8, and combined with the initial elution supernatant. The combined elution mixture was reduced and alkylated via the addition of 4 mM DTT and 10 mM iodoacetamide for 1 h at 37°C in the dark. An additional 0.5 μg Trypsin Gold was added to the samples, and the digestion was allowed to proceed overnight at 37 °C. Samples were acidified with TFA and cleaned up via MCX stage strip. We dried the post-cleanup samples via speed vacuum.

We reconstituted the dried peptide fractions in 97.9:2:0.1 H_2_O: acetonitrile (ACN): formic acid (FA) (load solvent) and analyzed ∼400 nanograms by capillary LC-MS with a Thermo Fisher Scientific, Inc (Waltham, MA) Dionex UltiMate 3000 RSLCnano system online with an Orbitrap Fusion mass spectrometer (Thermo Scientific, Waltham MA). We injected peptides directly in load solvent and performed gradient separation on a self-packed C18 column (Dr. Maisch GmbH ReproSil-PUR 1.9 um 120 Å C18aq, 100 um ID x 40 cm length) at 55°C with the following profile: 5% B solvent from 0 – 2 minutes, 12% B at 5 minutes, 40% B at 150 minutes, 50% B at 180 minutes and 90% B at 182 minutes with a flowrate of 400 nl/min from 0 – 2 minutes and 315 nl/minute from 2.5 – 180 minutes, where solvent A was 0.1% formic acid in water and solvent B was 0.1% formic acid in ACN. We employed the following MS parameters: ESI voltage +2.1 kV, ion transfer tube 275°C; no internal calibration; Orbitrap MS1 scan 120k resolution (at 200 *m/z*) in profile mode from 380 – 1500 *m/z* with 50 msec injection time; 100% (4 x 10^5^) automatic gain control (AGC); high energy collisional dissociation (HCD) MS2 activation was triggered on precursors with 2 – 5 charges above 1x10^4^ counts; MIPS (monoisotopic peak determination) was set to Peptide; MS2 settings were: 1.6 Da quadrupole isolation window, 30% fixed collision energy, ion trap detection, 35 msec max injection time, 100% (1 x 10^4^) AGC, 45 sec dynamic exclusion duration with +/- 10 ppm mass tolerance.

### Statistical Analysis

Values were expressed as the mean ± SEM. In comparisons made between two groups, Student’s *t*-tests were performed using GraphPad Prism (San Diego, CA). For comparisons among more than two groups, we used analysis of variance (ANOVA) with a Newman-Keuls post hoc test. Values with p<0.05 were considered statistically significant.

**Supplemental Figure 1 (Related to Figure 1).**

A) OCT frozen tissue sections stained with Oil Red O from WT mice treated with and without LPS. B) Quantification of LDs from mice treated with and without LPS; n=4 three images per section. ****<0.001. C) Liver TG was determined via colorimetric assay from mice treated with and without LPS; n=4. **<0.01. D) Liver fatty acid oxidation rates were determined as described in ^59^ from mice treated with and without LPS; n=4. **<0.01. E) Serum β-hydroxybutyrate was quantified from mice treated with and without LPS; n=4. ***<0.005. F) PGE_2_ was quantified from mice treated with and without LPS; n=6. G) Western blot analysis of cellular fractions. PLIN2-LD, αTubulin-Cytosolic, SERCA2-ER, COXIV-mitochondria, H3-nuclear. H) Western blot analysis of LDs isolated from mice treated with and without LPS. I) Quantification of Western blots normalized to total protein loading.

**Supplemental Figure 2 (Related to Figure 1).**

A) Western blot analysis of tissues collected from control WT and PLIN2 KO mice; n=3. Western blot analysis of PLIN2 was performed on liver, kidney, heart, eWAT, and muscle to confirm PLIN2 ablation. B-P) Innate immune, cytokine, and oxylipin gene expression in livers of mice treated with Veh or LPS. Data represent the mean ± SEM. Statistical significance was determined by two-way ANOVA. n= 6-8 mice per group. *P<0.05, **P<0.01, ***P<0.005, ****P<0.001.

**Supplemental Figure 3. (Related to Figure 1)**

A) WT and PLIN2 KO liver lysates from mice treated with and without LPS were harvested and clarified. Cytokine levels were assessed using the Proteome Profiler mouse Cytokine-XL Array Kit (R&D Systems). Membranes were imaged and quantified via densitometry. B) Results of cytokines impacted by PLIN2 ablation are shown as a fold change relative to internal controls. All antibodies are spotted in duplicate. Liver lysates from four mice per treatment group were combined 1:1:1:1 for a total of 400 μg total protein. Statistically significant changes were determined by two-way ANOVA. *P<0.05, **P<0.01, ***P<0.005, ***P<0.001.

**Supplemental Figure 4. (Related to Figure 2)**

A) Confocal images of AML12 cells treated with Veh or LPS and counterstained with AutoDOT to visualize LDs. B) Quantification of the number of LDs per cell. Quantification of AML12 LD numbers was performed on 20-40 cells; *P<0.05 as determined by Student’s t-test. C) cPLA2 activity was determined from AML12 cell lysates as described in the Methods section; n=8. ***P<0.001 as determined by Student’s t-test. D) Prostaglandin E2 levels were determined from AML12 cell lysates treated with Veh or LPS for 16 h; n=16. **P<0.01 as determined by Student’s t-test. E-G) Similar measures as A-D; only the data were collected from primary hepatocytes isolated from C57BL/6 mice; n=6-16. *P<0.05, **P<0.01 as determined by Student’s t-test. H) Primary mouse hepatocytes were isolated and treated with LPS medium. We performed the EicosaCell assay to visualize PGE2 as described in the Methods section. Confocal imaging of Cy3-labeled PGE_2_ and Cy5-Plin2-labeled cells revealed sites of synthesis in the cell. I) Prostaglandin E2 levels were determined from primary mouse hepatocytes isolated from control and PLIN2 KO mice treated with Veh or LPS; n=12. ****P<0.001 as determined by two-way ANOVA. J-K) Gene expression of IL6 (J) and COX2 (K) from primary mouse hepatocytes isolated from control and PLIN2 KO mice treated with Veh or LPS; n=6. ****P<0.001 as determined by two-way ANOVA.

**Supplemental Figure 5. (Related to Figure 2)**

Overview of arachidonic acid, linoleic acid, DHA, and EPA metabolites produced by lipoxygenases (LOX). A-G) Quantification of HETEs; H-I) HODEs; J-K) oxoODEs; L) LTB4; M) 12-oxoLTB4; N-T) HEPEs; and U-W) DiHETEs from hepatic tissue lysates of control and PLIN2 KO mice treated with Veh or LPS; n=4-6. ns indicates no statistical significance, *P<0.5, **P<0.01, ***P<0.005, ****P<0.001 as determined by two-way ANOVA.

**Supplemental Figure 6. (Related to Figure 2)**

Overview of arachidonic acid, linoleic acid, DHA, and EPA metabolites produced by cytochrome P450s (CYP). A-D) Quantification of EETs; E-H) DiHETs; I-J) EpOMEs; K-L) DiHOMEs; M-N) TriHOMEs; and O-Q) oxoETEs from hepatic tissue lysates of control and PLIN2 KO mice treated with Veh or LPS; n=4-6. ns indicates no statistical significance, *P<0.5, **P<0.01, ***P<0.005, ****P<0.001 as determined by two-way ANOVA.

**Supplemental Figure 7. (Related to Figure 3)**

A) The APEX2 biotin ligase experimental design. B) The APEX2-fusion constructs designed to determine the PLIN2-interactome. C) Western blot of cell lysates of mouse embryonic fibroblasts transduced with lentiviral particles containing APEX2-fusion constructs outlined in B. Cells were treated with LPS overnight, loaded with biotin-phenol, and APEX2 was triggered with H_2_O_2_ as described in the methods section. Samples were probed with anti-biotin to confirm that the APEX2 biotin ligase was working in each of the cell line constructs. D) The biotin-ligase experimental design used to determine the PLIN2-interactome. E) Summary of the number of peptides and proteins identified from each of the construct samples. F) Venn diagram showing the number of protein IDs that overlapped and that were distinct in each of the construct samples.

G) K-means statistical analysis was used to generate heatmap clustering of protein IDs into ten distinct expression patterns of proteins identified from the APEX2-PLIN2 samples. Yellow indicates ion abundance increased due to LPS treatment, while blue indicates decreased ion abundance. Protein IDs were mapped to specific pathways using PANTHER pathway analysis.

H) ShinyGO was used to group pathway enrichment, followed by I) KEGG-STRING analysis of the PLIN2 interactome. Yellow strings indicate a direct regulatory role of arachidonic acid metabolism on each pathway node.

**Supplemental Figure 8. (Related to Figure 3)**

A) Quantification of the quantitative immunoprecipitations shown in Figure 3G; n=3. *P<0.05 as determined by Student’s t-test. B) PLIN2 homology model^38^ based on the X-ray crystal structure of the PLIN3 C-terminus^84^ (PDB entry 1SZI), which was further refined using SwissModel and Phyre3D to define the PLIN2 lipid-binding pocket, with the N-terminus incorporated into the overall model. The C-terminal 4-helix bundle has been experimentally shown to contain the lipid-binding pocket^38^. Constraining ligand interactions to this region, we docked arachidonic acid to the PLIN2 homology model to define its binding capabilities. The top three arachidonic acid docking configurations are shown. B-C) These three models gave ΔG binding energies in the -20.4 to -7.2 kcal/mol range.

**Supplemental Figure 9. (Related to Figure 4-5)**

A) Ponceau S total protein loading for Western blot in Figure 4J. B) Quantification of acetylated lysine in liver lysates and isolated LDs from the Western blot in Figure 4J; n=3. ***P<0.005, ****P<0.001 statistically significant changes as determined by Student’s t-test. C) Immunohistochemistry in AML12 cells treated with vehicle or LPS overnight; red=anti-acetylated lysine, magenta=anti-PLIN2. Green overlay=colocalization analysis indicating regions of acetylated lysine overlapping with the PLIN2 signal, as determined by Pearson’s Correlation. Images show the full experiment with controls; select images are shown in Figure 4K. D) Quantification of PLIN2-acetylated lysine colocalization; n=20-40 cells. ***P<0.005 indicates statistically significant changes as determined by Student’s t-test. E) Ponceau S total protein loading for Western blot in Figure 5L and M. F) Quantification of phospho-tyrosine in liver lysates and isolated LDs from Western blot in Figure 5L. G) Quantification of phospho-serine in liver lysates and isolated LDs from Western blot in Figure 5M; n=3. *P<0.05 indicates statistically significant changes as determined by Student’s t-test.

---

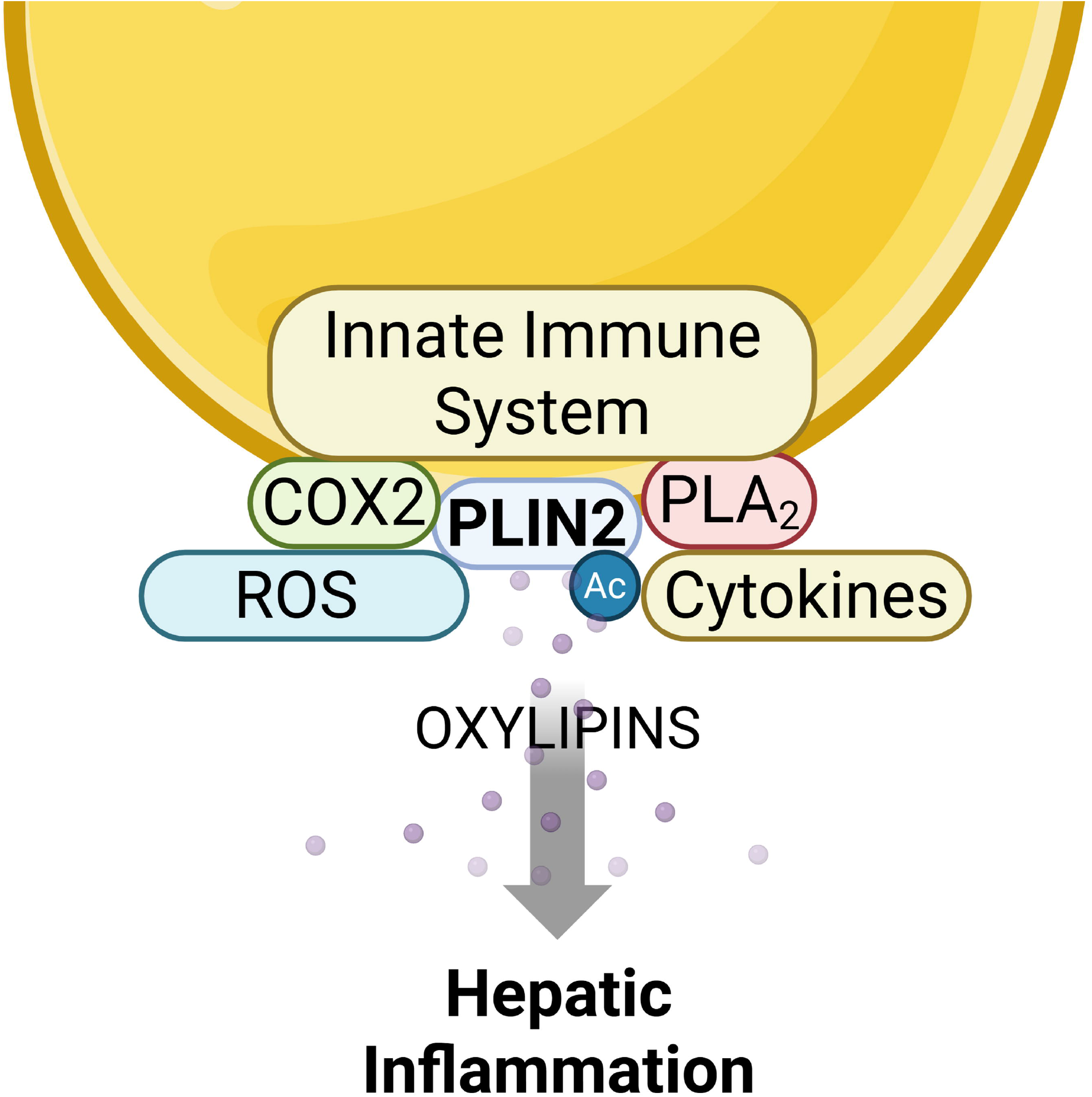

