## Supplemental Figures for "Acetylated PLIN2 coordinates the innate immune system and oxylipin synthesis on lipid droplets to propagate inflammation"

### Supplemental Figure 1

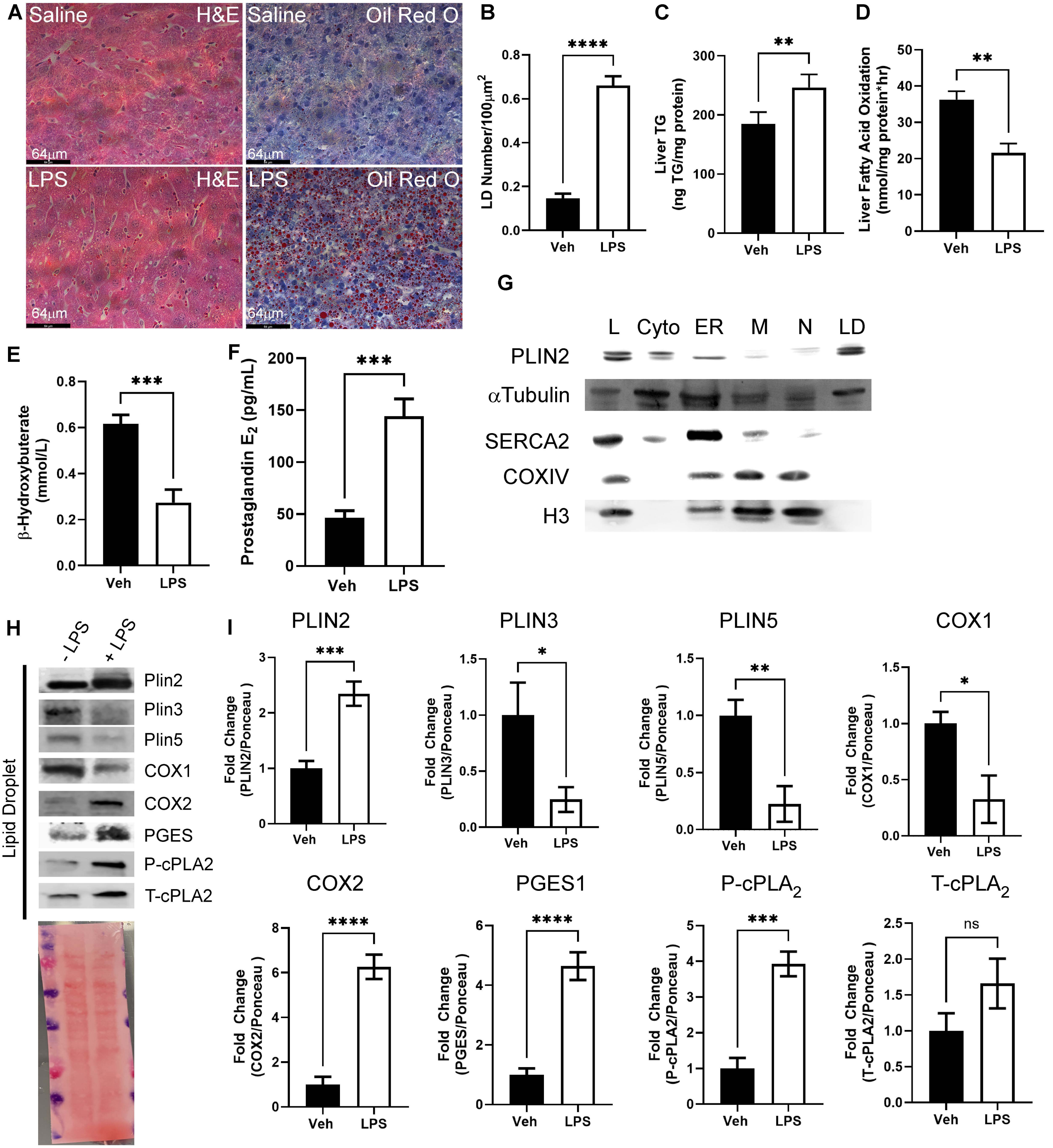

Supplemental Figure 2

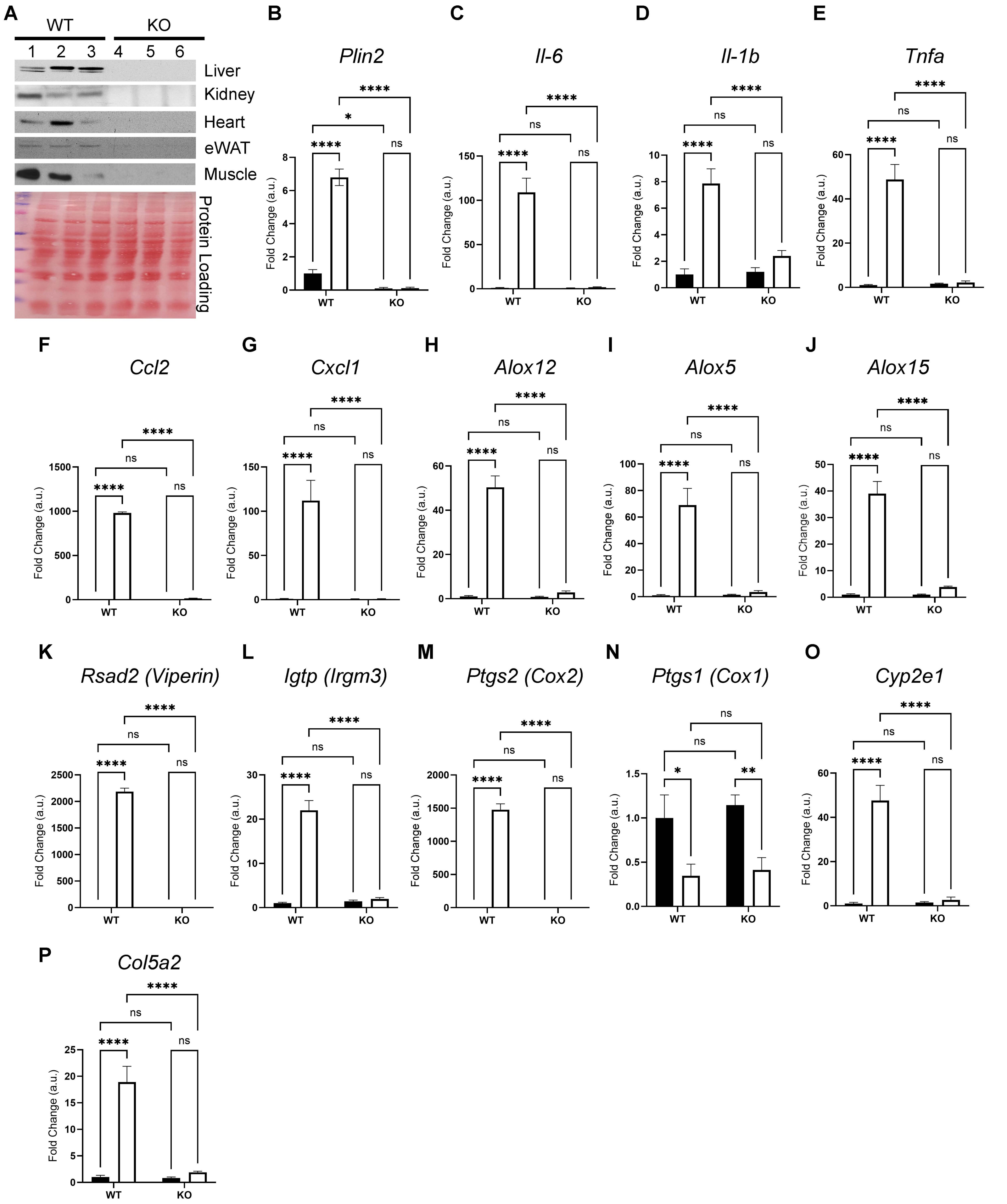

Supplemental Figure 3

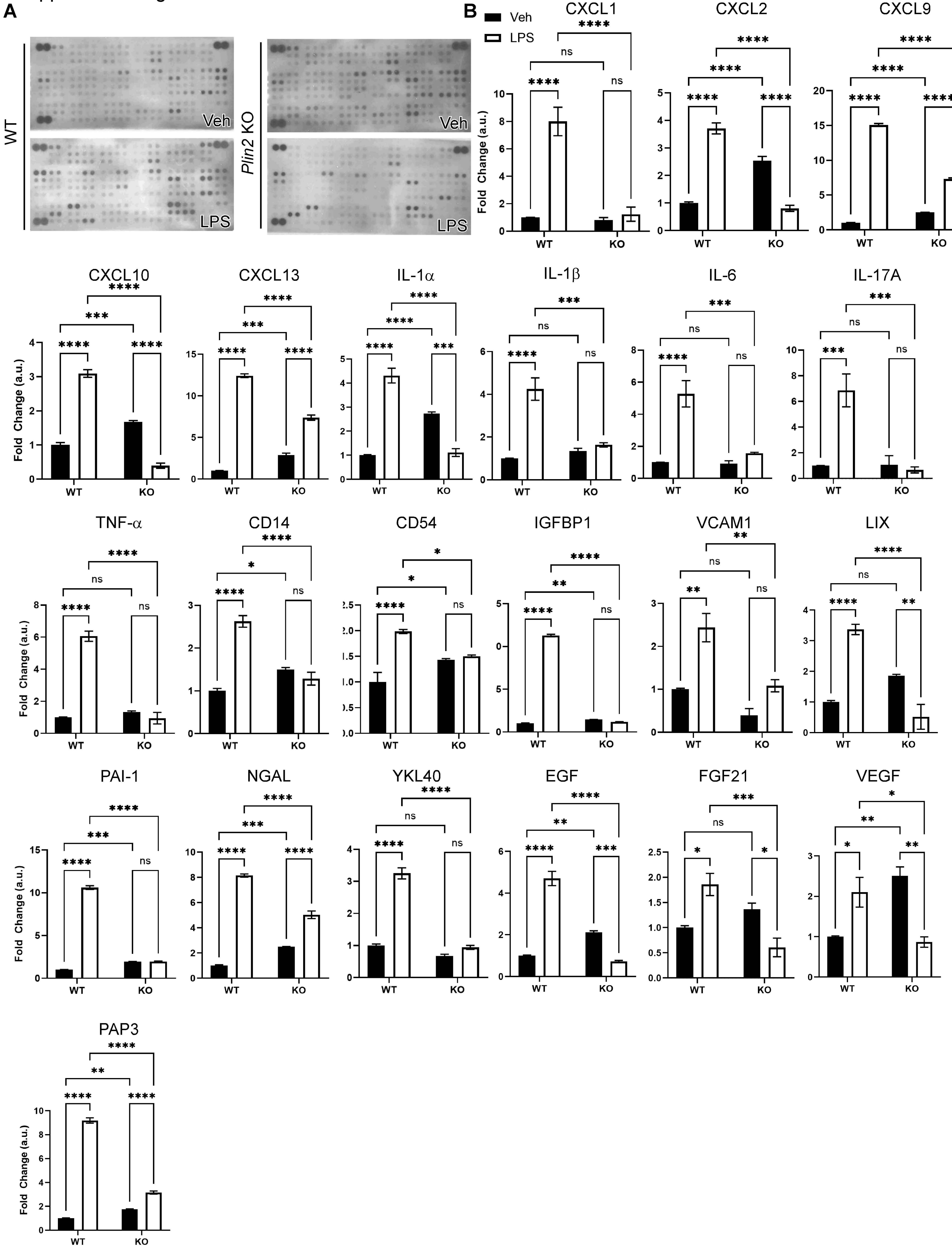

Supplemental Figure 4

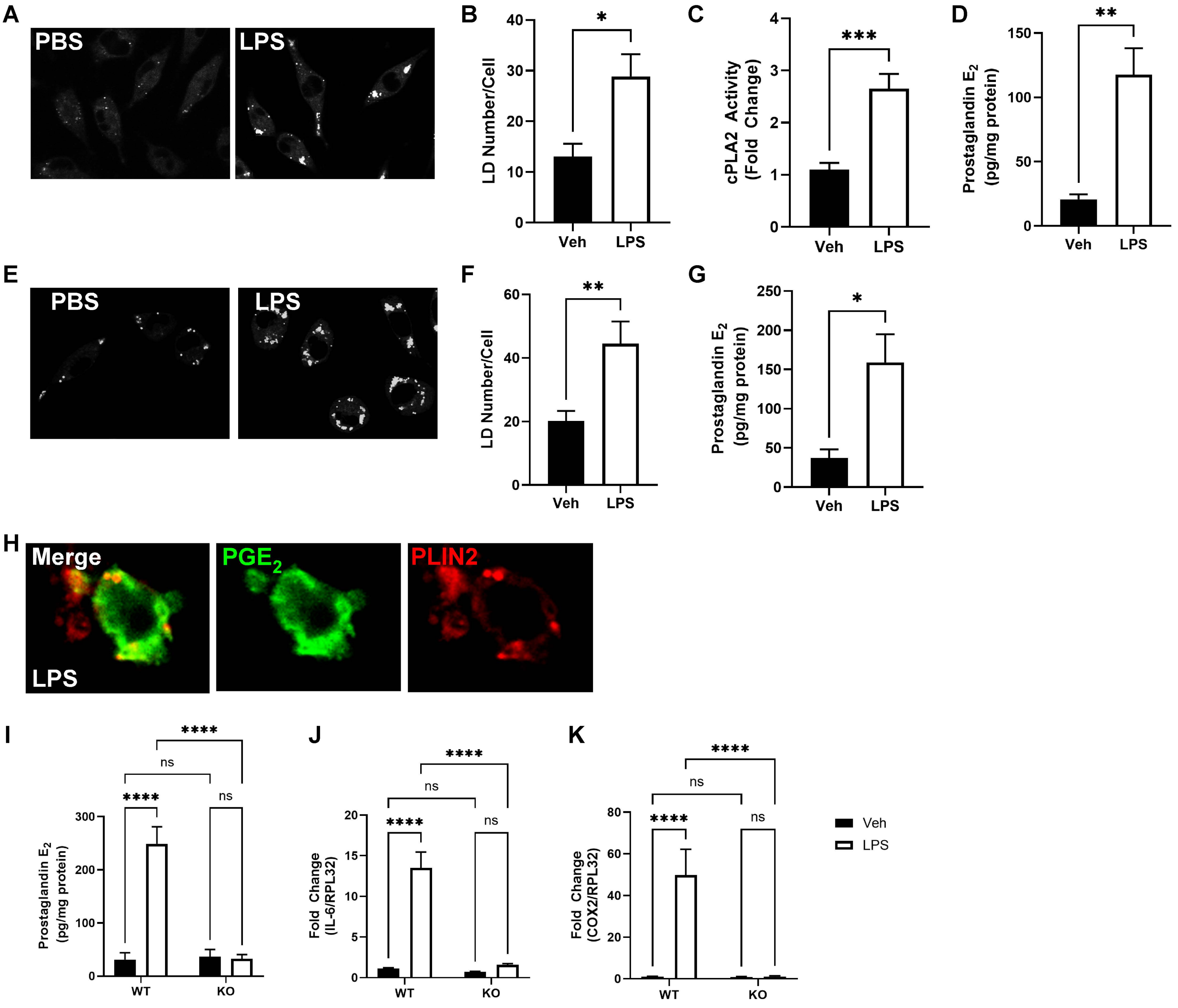

Supplemental Figure 5

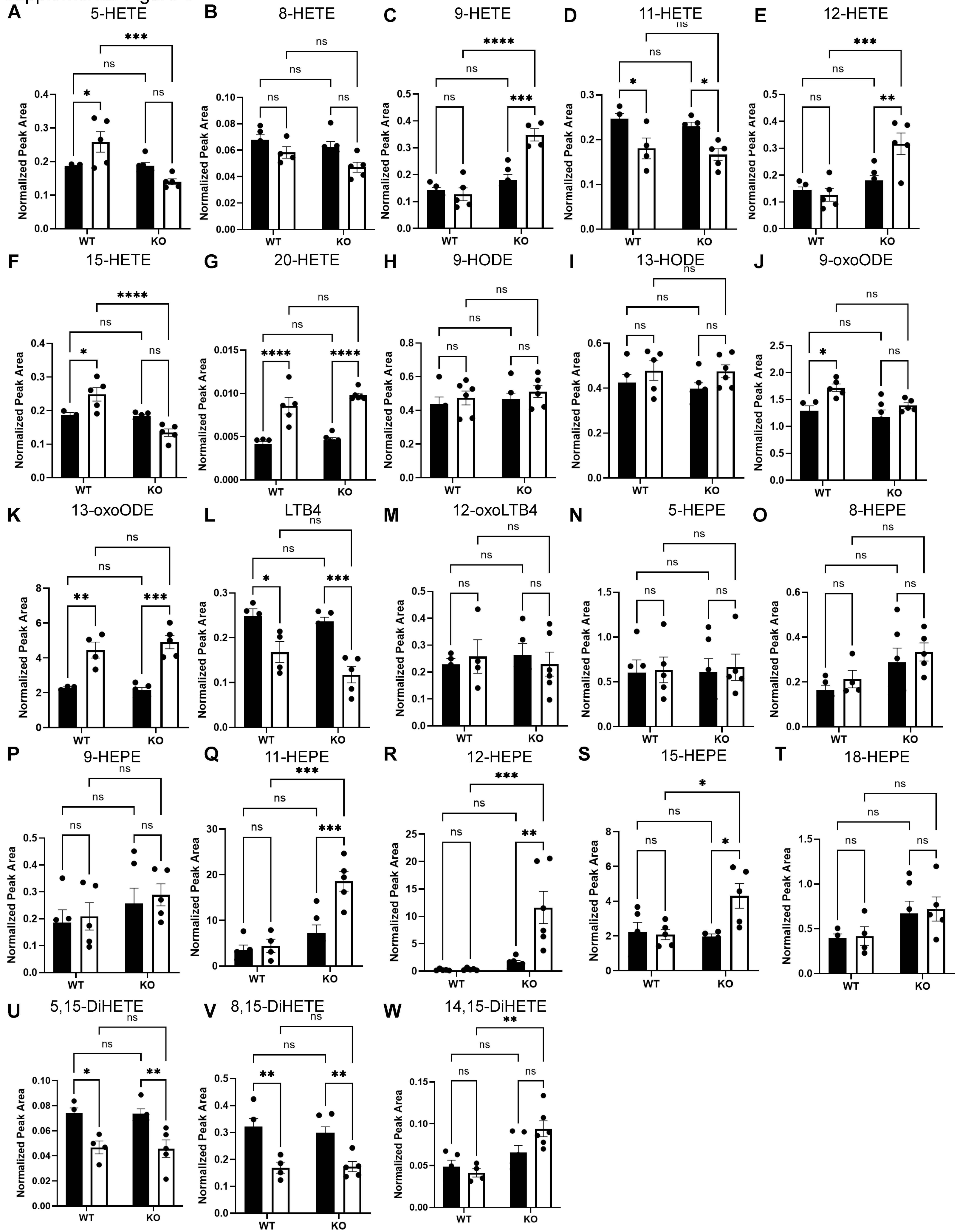

Supplemental Figure 6

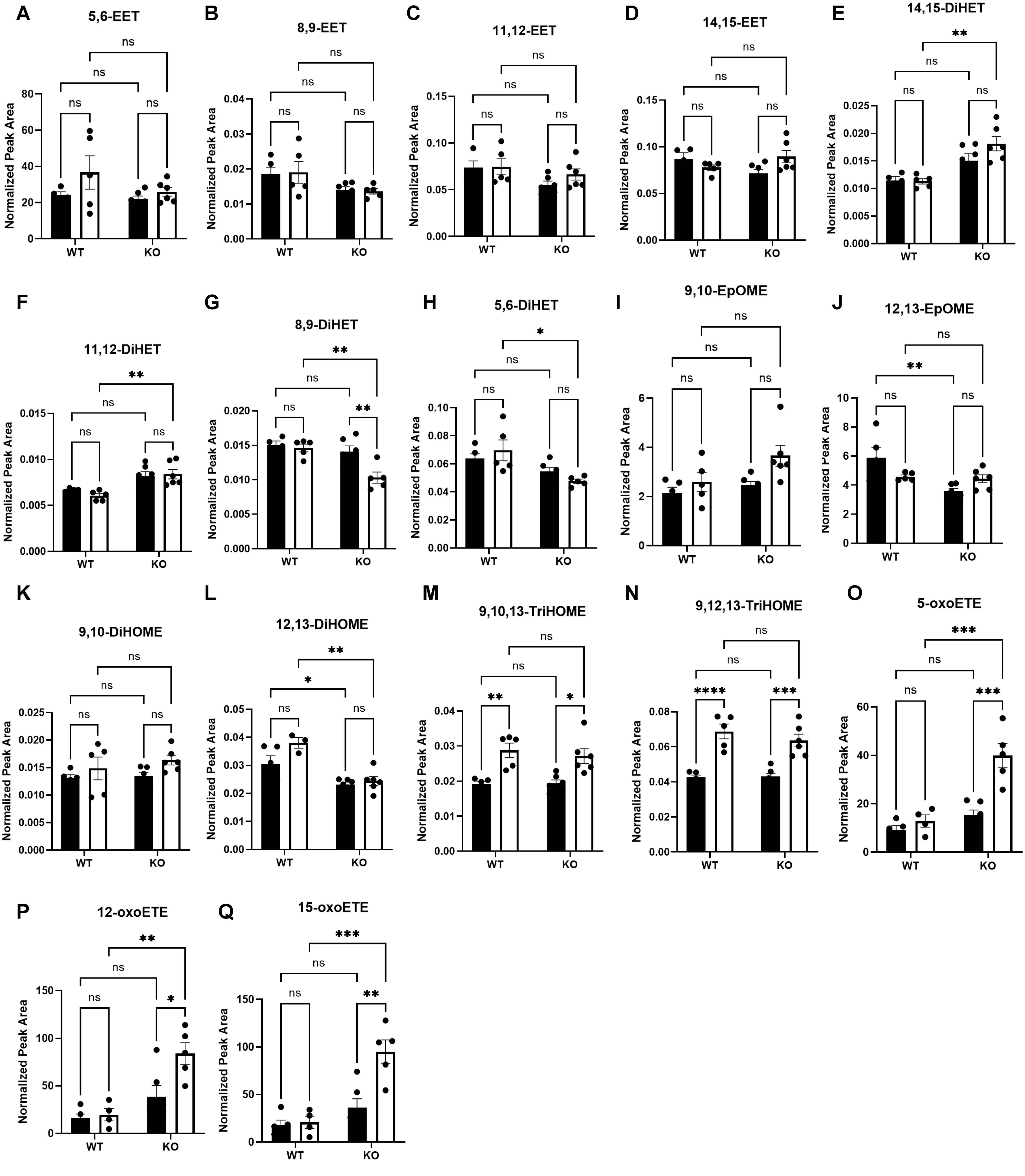

Supplemental Figure 7

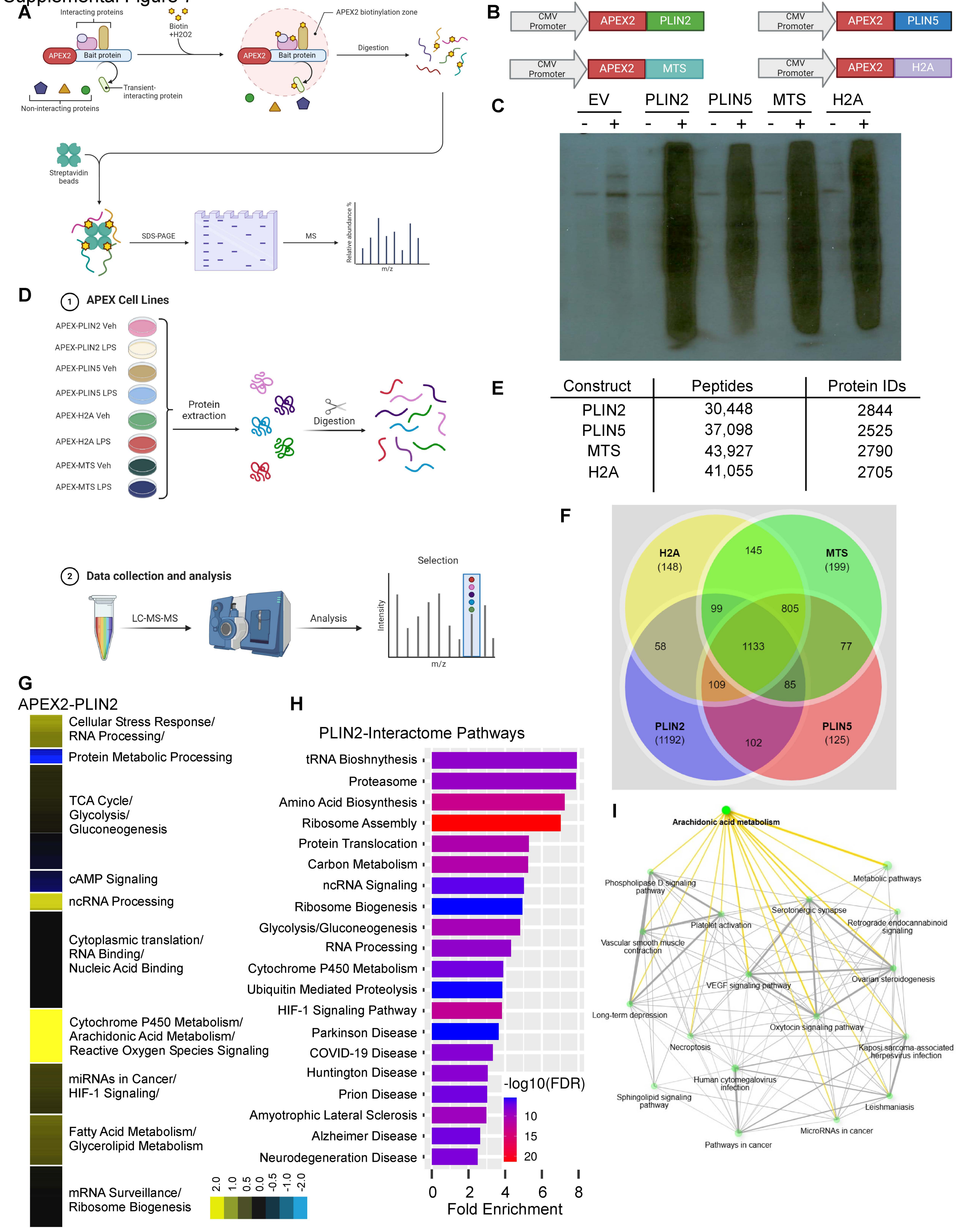


| Construct | Peptides | Protein IDs |
|---|---|---|
| PLIN2 | 30,448 | 2844 |
| PLIN5 | 37,098 | 2525 |
| MTS | 43,927 | 2790 |
| H2A | 41,055 | 2705 |

Supplemental Figure 8

A

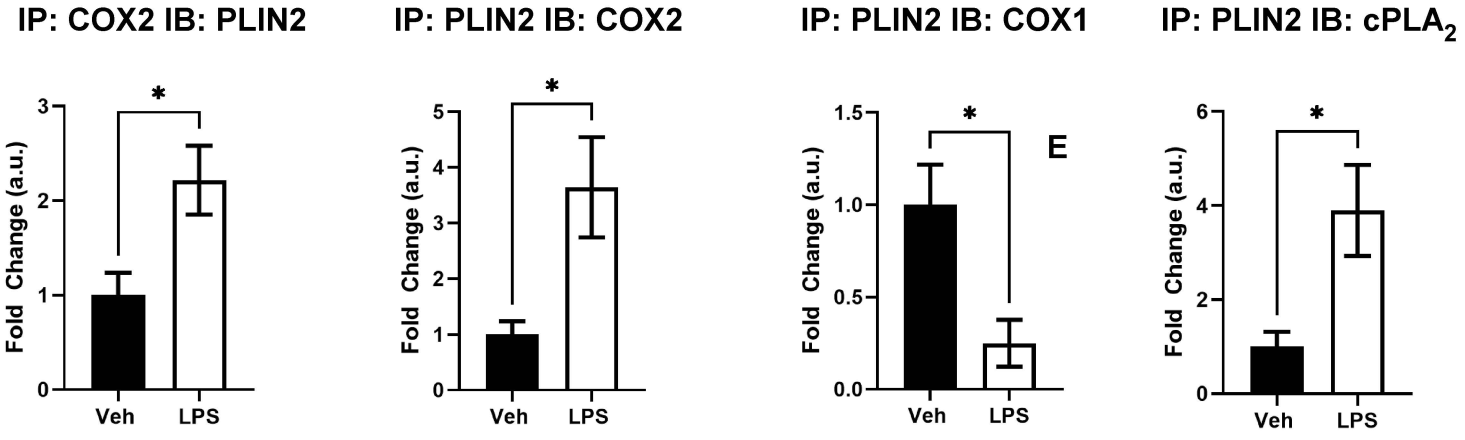


B

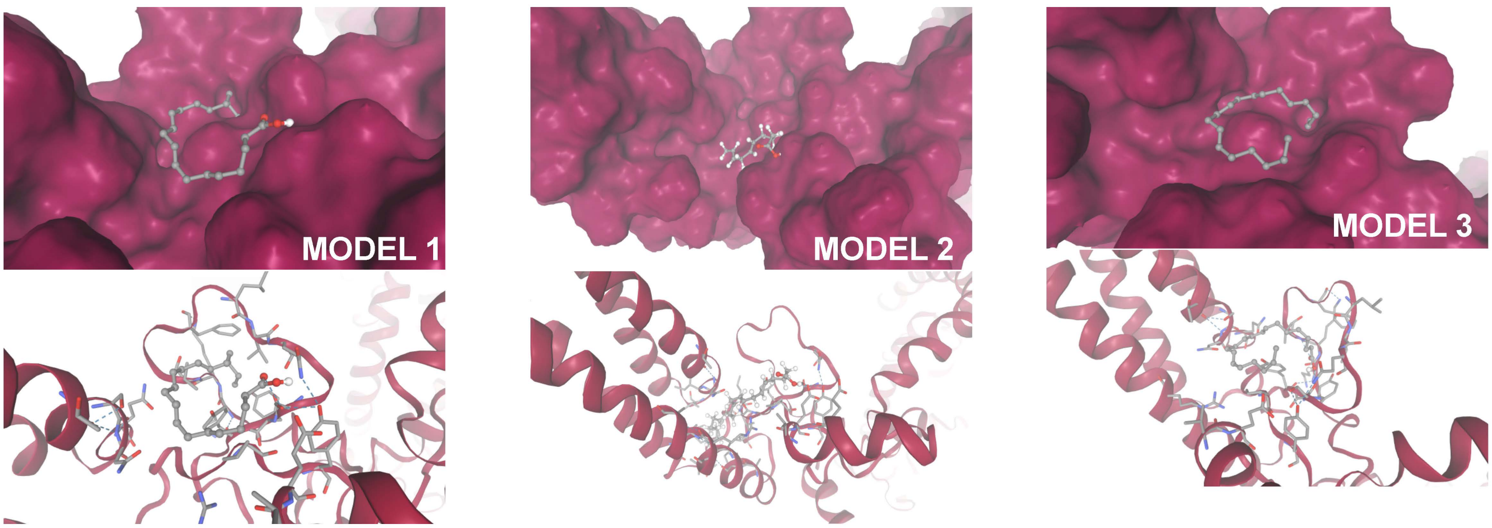


C

| Model | ΔG of binding |
|---|---|
| #1 | -20.4 |
| #2 | -13.6 |
| #3 | -7.2 |

Supplemental Figure 9

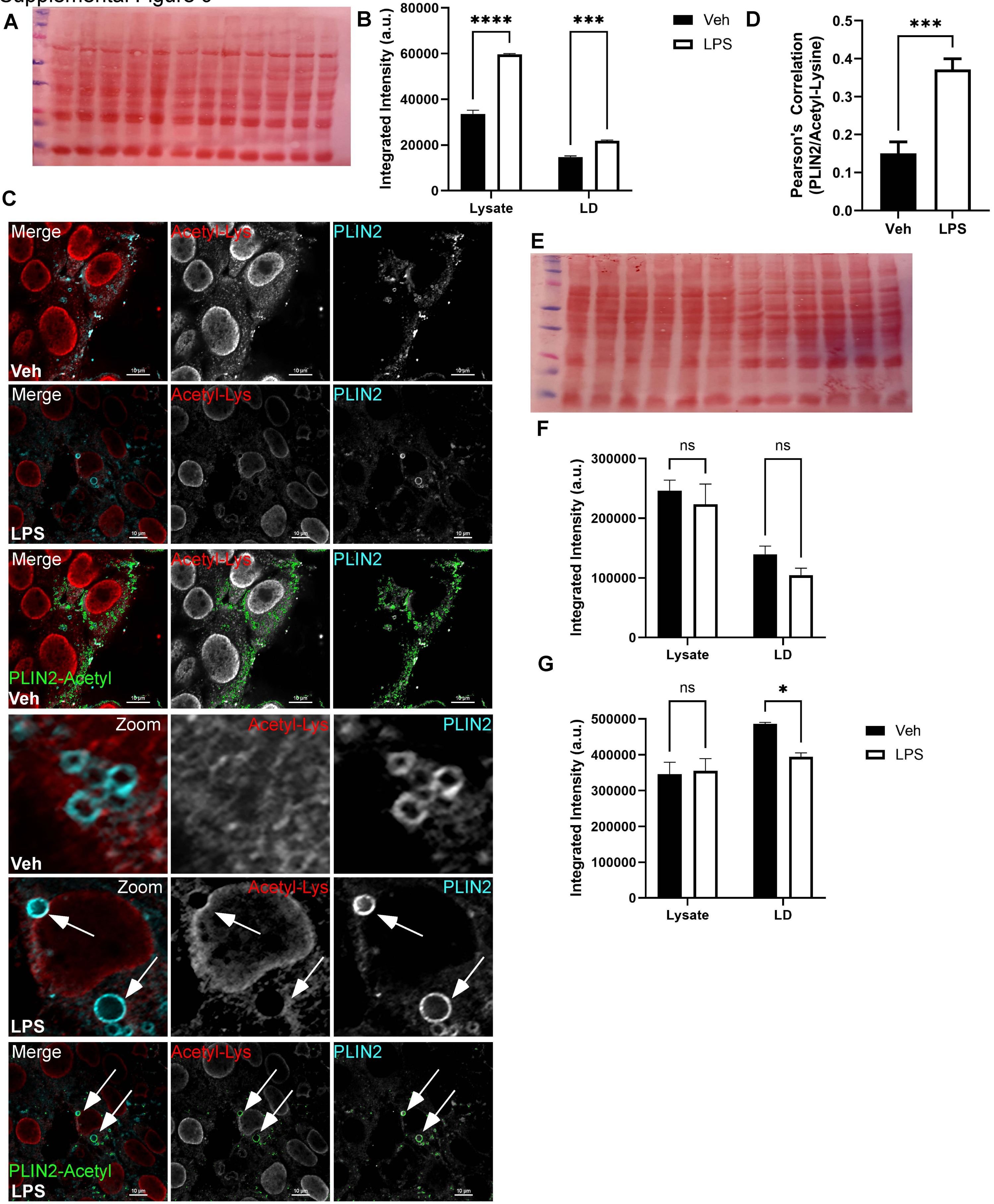
